# A systems genomics approach to uncover patient-specific pathogenic pathways and proteins in a complex disease

**DOI:** 10.1101/692269

**Authors:** Johanne Brooks, Dezso Modos, Padhmanand Sudhakar, David Fazekas, Azedine Zoufir, Orsolya Kapuy, Mate Szalay-Beko, Matthew Madgwick, Bram Verstockt, Lindsay Hall, Alastair Watson, Mark Tremelling, Miles Parkes, Severine Vermeire, Andreas Bender, Simon R. Carding, Tamas Korcsmaros

## Abstract

We describe a novel precision medicine workflow, the integrated single nucleotide polymorphism network platform (iSNP), designed to identify the exact mechanisms of how SNPs affect cellular regulatory networks, and how SNP co-occurrences contribute to disease pathogenesis in ulcerative colitis (UC). Using SNP profiles of 377 UC patients, we mapped the regulatory effects of the SNPs to a human signalling network containing protein-protein, miRNA-mRNA and transcription factor binding interactions. Unsupervised clustering algorithms grouped these patient-specific networks into four distinct clusters based on two large disease hubs, NFKB1 and PKCB. Pathway analysis identified the epigenetic modification as common and the T-cell specific responses as differing signalling pathways in the clusters. By integrating individual transcriptomes in active and quiescent disease setting to the patient networks, we validated the impact of non-coding SNPs. The iSNP approach identified regulatory effects of disease-associated non-coding SNPs, and identified how pathogenesis pathways are activated via different genetic modifications.

## Introduction

Precision medicine has been achieved in well demarcated monogenic diseases^1^ and in diseases where the pathogenic mechanism is well described such as the use of tamoxifen in HER2 positive breast cancer^2^. In diseases such as inflammatory bowel disease however, where there are multiple contributing factors to the disease pathogenesis, notwithstanding the complex genetics, precision medicine remains an aspiration. Therefore, complex integrative techniques are required to identify the individuals’ pathogenic disease pathways, to move towards a more precision medicine approach. With inflammatory bowel disease (IBD), the interlinked facets to disease are thought to be a dysfunction of the immune system in response to, as yet unclear, environmental triggers in a genetically susceptible host^3^. Focusing solely on genetic susceptibility, genome-wide association studies (GWAS) and subsequent fine-mapping of identified regions aimed to identify causal disease-associated variants^4,5^, but the clinical impact of these variants has yet to come to fruition. The functional annotation of SNPs in coding regions as an approach to define their biological impact has been utilised in obesity^6^, IBD^5^, and lung cancer^7^ allowing for computational workflows to prioritise SNPs for further analysis^8^. Understanding the function of SNPs in non-coding regions of the DNA, however, remains challenging and even the most refined fine-mapping identifies disease causing SNPs in areas that have yet to be annotated^5^. In a type of IBD, called ulcerative colitis (UC), coding SNPs (found in exonic regions) that alter amino acid composition and the function of the translated proteins comprise less than 10% of the total UC associated SNPs^9^. These coding SNPs do not cause the expected impaired intestinal barrier function or inflammation as a pathognomic features of ulcerative colitis^10^. Identifying the functional attributes of the remaining 90% SNPs located in non-coding regions would expand the utility of complex disease-associated SNPs. Analysing these non-coding SNPs allows the identification of novel pathogenic pathways, and potentially patient-specific disease susceptibility, and thus enabling precision therapy.

For functional annotation of SNPs in non-coding regions, a key question is whether the SNPs affect gene expression. The ways in which a SNP can regulate gene expression include affecting long non-coding RNAs, splicing, or transcription factor binding sites (TFBS) in enhancer regions and within introns^11^. A further way for a SNP to affect gene regulation is by affecting miRNAs which modulate gene expression at the post-transcriptional level by reducing mRNA half-life and stability. miRNAs bind to their complementary recognition sequence, a miRNA-TS on the mRNA, thereby targeting the mRNA for destruction. Functional miRNA-TS-s have been found in open reading frames including exonic and intronic regions as well as in the 5’ untranslated regions^12–15^. There are a multitude of predictive algorithms for the identification of splicing enhancer or silencing sites^16–18^, or motifs for lncRNA binding^19–22^, however in this study we focused on two regulatory effects as examples; SNPs occurring in transcription factors binding sites and in miRNA target sites.

Individual disease-associated SNPs have been reported to affect TFBS and miRNA-TS in many diseases including diabetes, schizophrenia, coronary heart disease, and Crohn’s disease^23–26^. However, the combined regulatory effects of these non-coding SNPs have not yet been evaluated at a systems level. This is particularly pertinent in UC, which is a disease that reflects disturbances of complex intracellular and intercellular networks. A systems biology approach has been utilised with predictive network models that identified proteins involved in the pathogenesis of IBD in general^27^ but this approach was unable to take account of the regulatory effect of non-coding SNPs. To identify the effect of non-coding SNPs, we build on the concepts identified by Boyle et al (2017) to track the cumulative effects of multiple regulatory SNPs.

Using network biology approaches, that we have previously exploited to uncover novel important proteins in cancer biology,^28^ we aimed to further understand the pathogenic pathways of UC and to identify novel disease associated hidden proteins. These proteins are often hidden if one looks only conventional mutation and expression screens as they mostly act as direct interactors (first neighbours) of the proteins affected by the mutation. Similar studies have utilised the concept of first neighbour disease associated proteins in both diabetes^29^ and juvenile idiopathic arthritis^30^.

Consequently, we combined and refined systems genomics and network biology approaches into a novel workflow, named the integrative SNP Network Platform (iSNP), and demonstrated its applicability by analysing a UC-associated signalling network and by identifying patient clusters with distinct pathomechanisms contributing to UC. Within these clusters, we highlighted cluster-specific key players, some already supported in the literature. The iSNP approach also provided patient-specific pathogenic role for proteins whose contribution to UC pathogenesis was unknown. We then validated these predicted pathogenic effects using genotyped transcriptomics data. We show that integrating systems genomics and network biology data and analysis with machine learning approaches offers unique biological insights, and enables the scalable examination of patient-specific datasets for precision medicine.

## Results

### Constructing the UC-associated signalling network

To assess the regulatory effect of non-coding SNPs we first needed to reconstruct an interaction network around them containing the directly affected genes and those proteins that are indirectly affected by the SNP. We developed a novel workflow, the integrative SNP Network Platform (iSNP) to reconstruct such a network (Figure 1). To create a specific, UC-associated signalling network, we selected UC associated SNPs from publically available datasets; Jostins *et al*^9^ and the Broad Institute^31^ with published risk alleles that were finemapped on immunochip or had been finemapped by Fahr et al^31^. In circumstances where the SNPs were GWAS SNPs (not on immunochip), we only utilised them if the R^2^ value to a finemapped SNP was >0.8. To identify the effect of these SNPs in a patient-specific manner, using an East Anglian UK cohort of 377 patients from the UK IBD genetics consortium, we extracted SNP profile data from each individual patient. These patients had a total of 40 individual UC associated SNPs from which we identified 12 UC-associated regulatory SNPs localized within TFBS or miRNA-TS. We removed four SNPs that did not meet the stringent kinetic cutoffs for miRNA-TS or had a neutral response (the risk allele did not change miRNA-TS binding kinetics). The remaining eight SNPs were annotated to occur within 25 individual miRNA-TSs and four TFBSs (Table 1). Interestingly, three SNPs affecting PKCB, DMN3TB and HDAC7 are all annotated to the TFBS for the transcription factor SMARCA4. In Table 1, we summarised the known UC-associated information of the miRNAs and transcription factors that were affected by a SNP, and predicted the overall effect of the SNPs for each gene. Given that TFBS could be inhibitory or activatory (while miRNA-TS are generally considered as inhibitory), we manually curated the probable transcriptional response based on the literature (Table 1.).

**Table 1:**
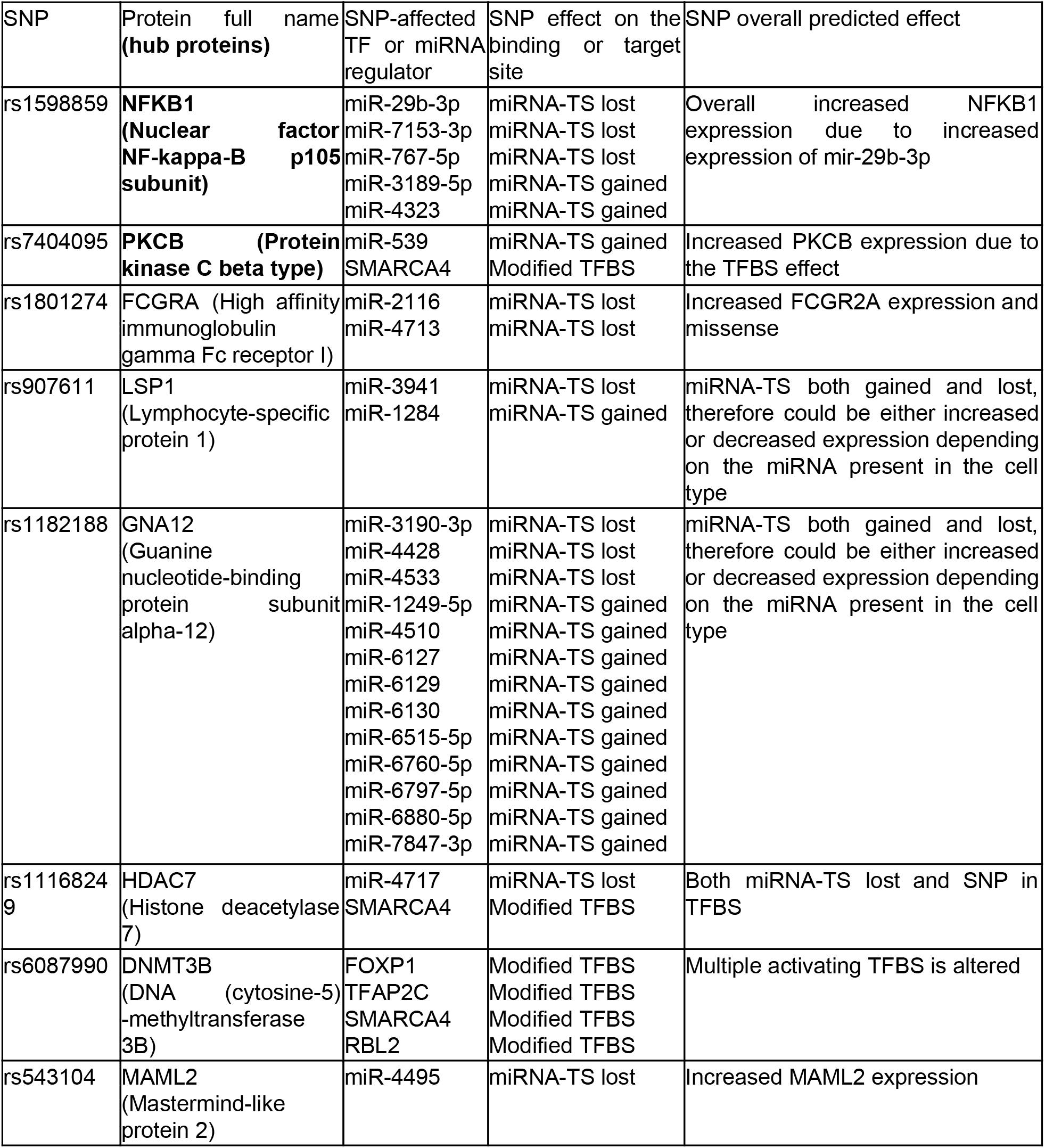
Ranked list of genes, miRNA-TS and TFBS affected by SNPs in the UC specific signalling network

**Figure 1.**
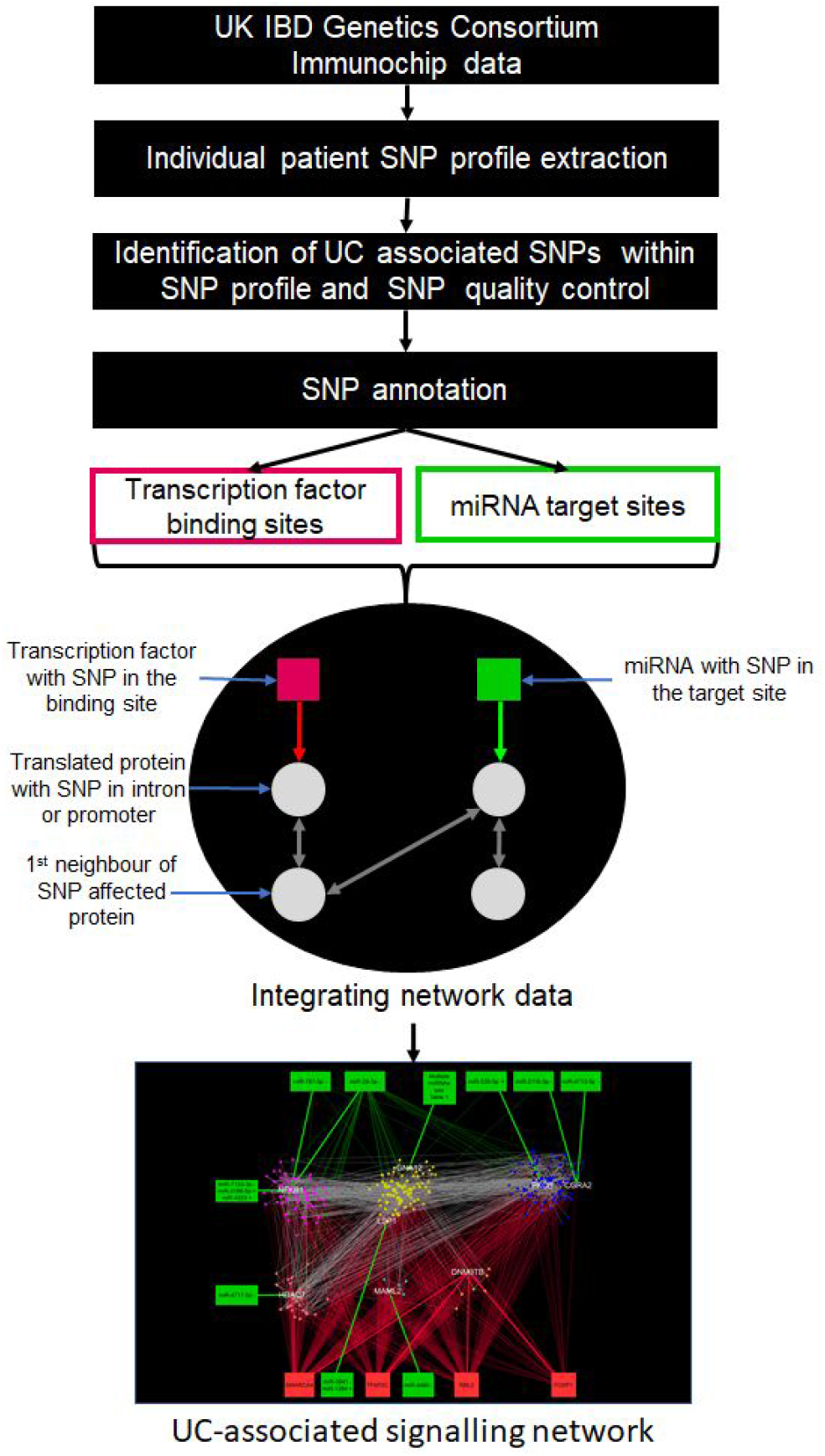
**The iSNP workflow** to reconstruct a disease specific network for non-coding SNPs. SNPs involved in patients were annotated based on that they occur within TFBS and miRNA-TS. Using regulatory interaction data sources we determined the potential affected proteins as well as their interaction partners from OmniPath, an integrated signalling network database.

The annotated SNP affected genes were translated to proteins, and using OmniPath^32^ (an integrated and comprehensive source for manually curated signalling interaction databases), we identified first neighbour interactors to the eight SNP affected proteins. Using Cytoscape^33^, we visualised the UC-associated signalling network containing protein-protein, miRNA and transcriptional interactions. In total, the UC-associated signalling network consisted of 247 protein nodes and 1,269 protein-protein interactions, regulated by 4 transcription factors and 25 miRNAs with altogether 682 regulatory interactions. The protein-protein interaction network was modularised for visualising the functions and key proteins of the network (Figure 2).

**Figure 2.**
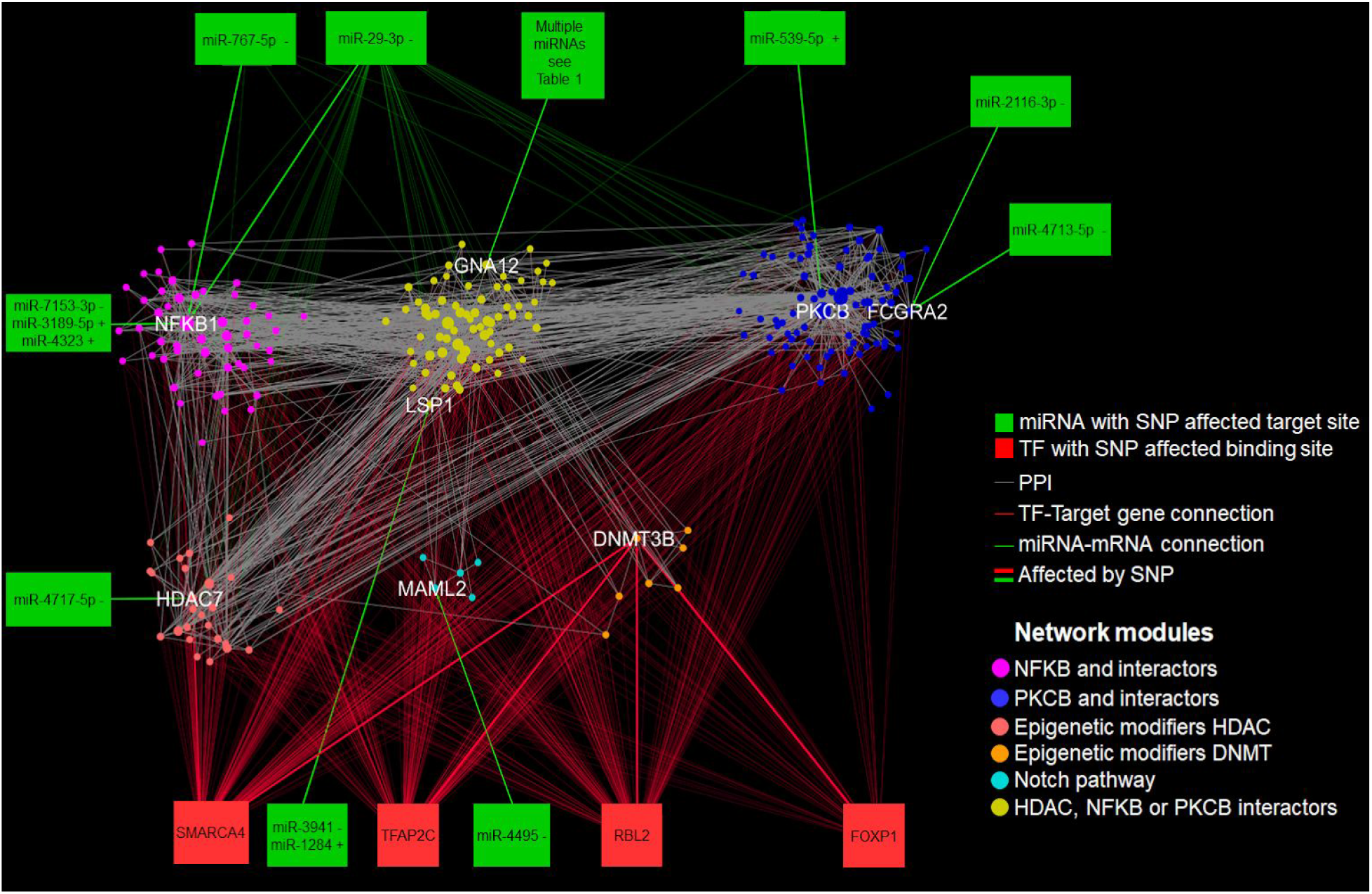
Visualization and modularisation of the UC-associated signalling network. The UC-associated signalling network containing proteins affected by UC associated SNPs, their interactor partners as well as the TFs and miRNAs whose binding site or target site are affected by a SNP. Circles represent proteins and squares represent regulators (TFs: red and miRNAs: green). The bold lines (edges) are the SNP-affected miRNA or TFBS, with the narrow lines (edges) representing known regulatory or protein-protein interactions. Nodes are coloured according to network modules and the name of SNP-affected proteins are in white. The miRNAs with a “+” symbol have putatively gained target site and miRNAs with a “−“ symbol have putatively lost their target site in UC patients.

In the UC-associated signalling network, the two central hub proteins (the proteins with the greatest numbers of connections or interactors) were NFKB1 (Nuclear Factor Kappa B Subunit 1) and PKCB (Protein Kinase C Beta). NFKB1 is one of the key regulators of the chronic mucosal inflammation driven by activated effector immune cells, which produce pro-inflammatory cytokines such as tumour necrosis factor-alpha and interleukin-6^34^. Protein Kinase C has been implicated in the pathogenesis of inflammatory bowel disease via effects on the colonic mucous layer^35^, colonic microbiota^36^ and cell junctions^37,38^, therefore both hub proteins are known to be involved in UC^39,40^ and were therefore expected to emerge from this analysis validating the iSNP method. The remaining six SNP-affected proteins were termed as non-hub SNP-affected proteins due to their lower number of interactions.

When analysed in more detail, the UC-associated signalling network consisted of six distinct but intertwined network modules. Each module is centred around a key signalling protein directly affected by a SNP (Figure 2). The three most abundant modules are formed mainly by the interactors of 1) PKCB and FCGR2A (Immunoglobulin G Fc Receptor II) (88 proteins), 2) NFKB1 (51 proteins), and 3) the binding partners of LSP1 (Lymphocyte Specific Protein 1) and GNA12 (Guanine Nucleotide-Binding Protein Subunit Alpha-12) that contained interactors of both NFKB1 and PKCB (71 proteins). We also identified two epigenetic modules around 4) Histone Deacetylase 7 (HDAC7; 25 proteins), and 5) around DNA Methyltransferase 3 Beta (DNMT3B; 7 proteins). These two epigenetic regulators are affected by SNPs altering not only miRNA-based post-transcriptional regulation (as in the other modules), but also transcriptional regulation (Table 1). Lastly, 6) a module containing MAML2 and members of the Notch pathway was identified.

### Identification of patient-specific clusters based on the networks of affected proteins

We then investigated how the UC-associated signalling network differed in each of the 377 UC patients. Based on the set of SNPs present in each patient, we defined patient-specific UC-associated signalling networks, called ‘network footprints’. The network footprint of each patient contained the proteins encoded by the SNP-affected genes and the interactors of these proteins, *i.e.* their first neighbour proteins^28^. Unsupervised hierarchical clustering using different linkage algorithms and multidimensional scaling of the network footprints of 377 patients stratified the patients into the same four distinct clusters (Figures 3a and 3b). For the distribution of patients in the four clusters, see Supplementary Table 2.

**Figure 3.**
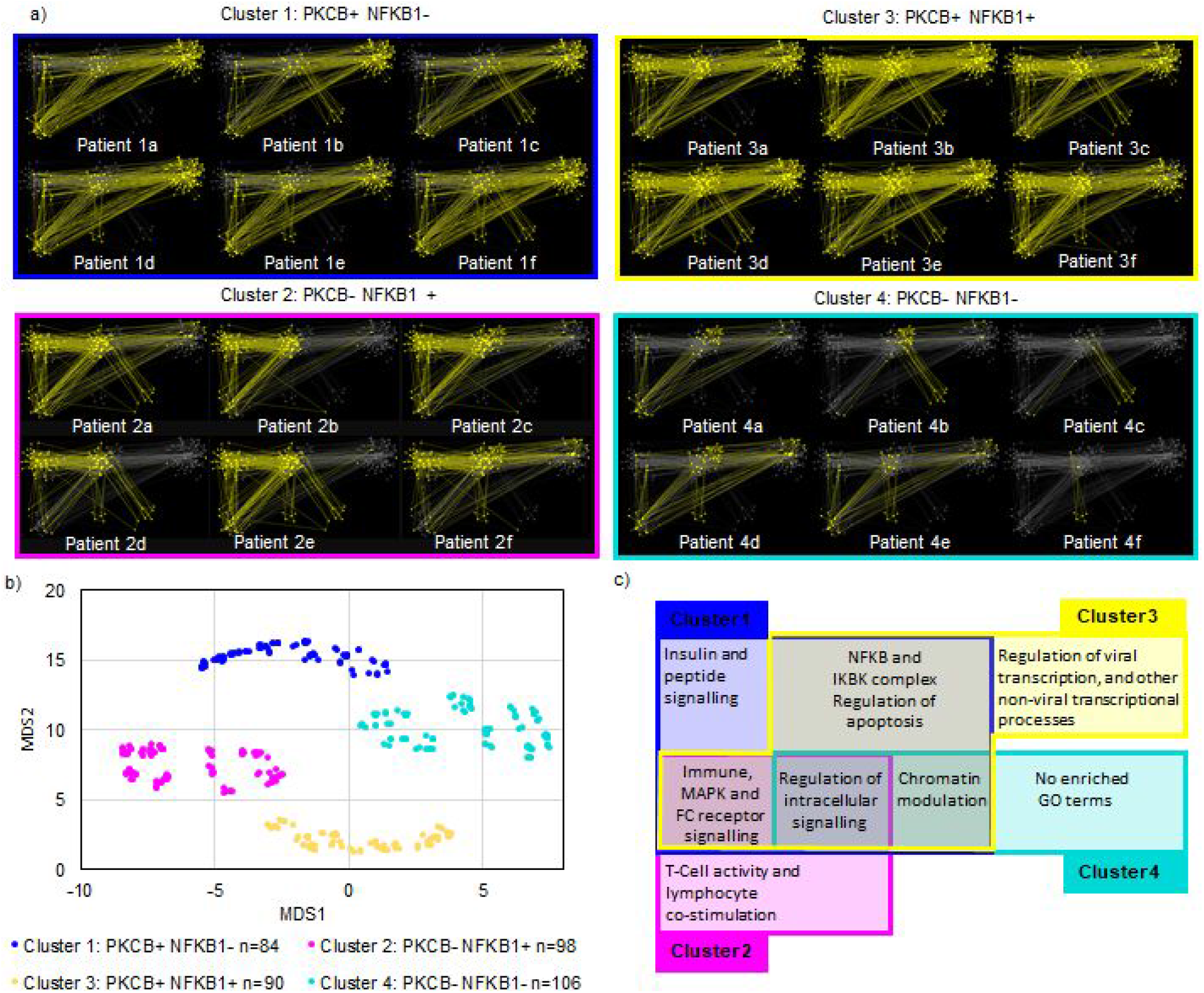
Unsupervised clustering of the patients and gene ontology of the clusters. a) Examples of patient-specific network footprints. b) Visualising of the clustering *via* multidimensional scaling c) Venn diagram of overrepresented Gene Ontology terms found in >50% of patients from each cluster. Complete GO data can be found in Supplementary Table 3. The “+” or “−” symbol means that in a given cluster the hub protein is either present or absent from those network footprints.

The first cluster contained the network footprints of patients whose SNPs were related to PKCB (denoted PKCB+, NFKB1− ; Figure 3 top left patient examples 1a-f), with the second cluster containing network footprints of patients with SNPs related to NFKB1 (PKCB− NFKB1 + ; Figure 3 bottom left patient examples 2a-f). In the third cluster, the network footprints contained both PKCB and NFKB1 SNPs (PKCB+ NFKB1+ ; Figure 3a top right patient examples 3a-f) while the network footprints of the fourth cluster had neither PKCB nor NFKB1 affected (PCKB− NFKB1− ; Figure 3 bottom right patient examples 4a-f).

To further characterise the different pathogenic pathways in the UC-associated signalling network (Figure 2), we conducted a patient cluster specific Gene Ontology (GO) enrichment analysis (see Supplementary Table 3 for full details). Altogether, we found 645 GO terms that were enriched in at least one patient. Next, we analysed which GO terms were enriched in more than 50% of patients in a given cluster. This led to two GO Biological Processes, which were represented in all four clusters: “Regulation of intracellular signal transduction” and “Positive regulation of response to stimulus”, confirming that intracellular signalling is a major player in UC pathogenesis. This annotation was statistically confirmed by the use of the whole OmniPath database as a background for the enrichment analysis indicating that even considering a large signalling network, UC affected processes are concentrating around regulatory functions of the signalling process^41^. Immune system pathways, such as the Fc receptor signalling pathway, immune response-regulating signalling pathway, were common to clusters with NFKB1 and PKCB hubs (Clusters 1-3). Chromatin modulation was a common GO Biological Process in Clusters 1, 3 and 4 suggesting the importance of epigenetic functions in these clusters (Figure 3c).

Three of the four clusters had cluster specific GO Biological Process terms. Response to insulin and peptide signalling was specific to Cluster 1 (PKCB+ NFKB1−). In Cluster 2 (PKCB− NFKB1+) the GO Biological Processes T-cell co-stimulation were enriched. Viral transcription and non-viral transcriptional Biological Processes were specific to Cluster 3 (PKCB+ NFKB1+). Meanwhile, Cluster 4 (PKCB− NFKB1−) did not show enriched GO terms specific to its patients. This result suggests a higher heterogeneity in Cluster 4, and necessitated a more detailed analysis to identify the key contributors of the UC pathogenesis here as it cannot be explained by current known mechanisms contributing to UC. To do this, we focused on the non-hub SNP-affected proteins we identified earlier (Figure 2), LSP1, MAML2, DNMT3B and HDAC7; the key proteins in Cluster 4.

### Non-hub SNP-affected proteins impact inflammation regulation

Given that NFKB1 targets are markers for increased inflammation in UC^42^, we aimed to analyze whether the presence (or absence) of non-hub SNPs were associated with up or downregulation of NFKB1 target genes in colonic biopsies. For this, we analysed transcriptomic data from paired inflamed and non-inflamed colonic biopsies from 44 UC patients with defined genetic backgrounds to capture differences in the expression of inflammatory genes. These patients were from the IBD referral centre in Leuven, Belgium, all with severe disease necessitating the use of anti-Tumour Necrosis Factor antibodies. Using the same iSNP workflow as described above for the UK IBD genetics consortium cohort, we confirmed that the SNP profiles from this independent Leuven patient cohort were similar to the UK IBD cohort. The only exception was that the Leuven patient cohort had an additional 25 UC associated SNPs indicating a higher coverage of known UC associated SNPs (Supplementary table 5). We reconstructed the patient network footprints and clustered the Leuven patient cohort. These clusters recapitulated the four clusters from the UK IBD cohort confirming that even with higher SNP coverage, UC patients can be grouped to these four clusters. We note that we also identified an additional hub protein, IL-7R affected by the SNPs rs11567701 and rs11567699 that were not present in the East Anglian cohort. This hub protein provided a higher granularity of the cluster structure, and its presence in the Leuven cohort resulted in an additional cluster. For the following validatory analysis, we focused only on the four patient clusters of the Leuven cohort that correlated with the four clusters of the UK IBD cohort.

We undertook Gene Set Enrichment Analysis focussing on 312 NFKB1 target genes in the transcriptomic data generated from all the patients in the Leuven cohort (Figure 4a). We investigated the SNP effect on the identified four non-hub proteins (LSP1, MAML2, DNMT3B and HDAC7) in all the four clusters l. (Figure 4b, p values in Supplementary Table 5).

**Figure 4.**
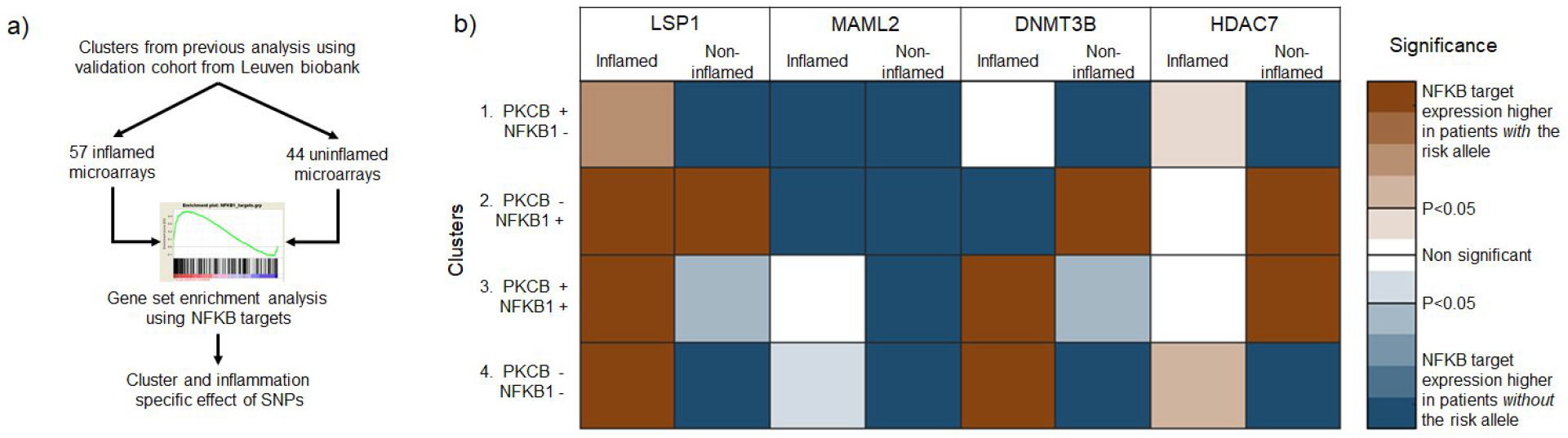
Non-hub SNP affected proteins have an inflammation specific effect on the expression of NFKB1 target genes. a) Analysis workflow for the transcriptomics based SNP validation; b) Heatmap of the Gene Set Enrichment Analysis (GSEA). The colours are representing whether the NFKB1 targets are over (brown) or under (blue) expressed regarding the presence of the SNP (for p values see Supplementary Table 6). The “+” or “−” symbols for each cluster mean that in a given cluster the hub protein is either present or absent from those network footprints

We found that the SNPs affecting the NFKB1 hub, or the PKCB hub are not consistently determining the changes in the NFKB1 target gene expression either in the inflamed or non-inflamed setting. This confirms that SNPs affecting non-hub proteins were playing a role in changes in NFKB1 targets gene expression (i.e., they contribute to the regulation of the inflammatory response), including in Cluster 4, where the NFKB1 and PKCB SNP risk alleles are not present.

### Non-hub SNP-affected proteins influence UC pathogenesis

To understand better how the four non-hub SNP-affected proteins (LSP1, MAML2, DNMT3B and HDAC7) can affect UC pathogenesis either in inflamed or non-inflamed settings, we evaluated their pathogenic or protective roles in each of the clusters.

LSP1 (Lymphocyte Specific Protein 1) is an actin binding protein involved in neutrophil and endothelial cell migration^43^. The risk allele rs907611 occurs within an LSP1 intron, and we annotated it to a loss and/or a gain of a miRNA-TS (Table 1). The presence of the risk allele at rs907611 was associated with decreased NFKB1 target gene expression in the non-inflamed cases in all clusters except Cluster 2 (PKCB− NFKB1+), where it was associated with a higher NFKB1 target gene expression. This suggested that in the non-inflamed setting rs907611 is protective. Contrary to this, in the inflamed setting the presence of the risk allele at rs907611, regardless of cluster, was associated with increased NFKB1 target gene expression, indicating that the effect of rs907611 is context specific, and may exacerbate existing inflammation by increasing NFKB1 target gene expression.

MAML2 (Mastermind Like Transcriptional Co-Activator 2) is a Notch pathway cofactor, which has a direct effect on NFKB1 translocation to the nucleus, thereby affecting downstream target gene expression^44–46^. The risk allele at rs543104 is annotated to lead to a loss of miR-4495 target site for *MAML2* mRNA that causes an increase in *MAML2* expression (Table 1). Interestingly, in the non-inflamed setting, the presence of the risk allele at rs543104 was associated with a decreased NFKB1 target gene expression in all the clusters. In the inflamed setting however, a different picture arose: the risk allele at rs543104 afforded protection (reduced NFKB1 gene target expression) only in Clusters 2 and 3 (where either NFKB1 or PKCB had a risk allele). When both hubs had risk alleles or neither of them (PKCB+ NFKB1+ or PKCB− NFKB1−), then the risk allele at rs543104 afforded no significant change in NFKB1 target gene expression.

DNMT3B (DNA Methyltransferase 3B) is a cytosine methyltransferase, which is involved in *de novo* DNA methylation and is essential for the establishment of DNA methylation patterns during development^47^. The risk allele rs6087990 occurs in the promoter region of DNMT3B, within a region of TFBSs enrichment for activating transcription factors such as FOXP1, TFAP2C and SMARCA4. In the non-inflamed setting, the presence of the rs6087990 risk allele was associated with reduced NFKB1 target gene expression only in the context of patients without the NFKB1 network hub being affected (Clusters 1 and 4). In Cluster 2 (PKCB− NFKB1+) patients having the risk allele at rs6087990 were associated with a significantly higher level of NFKB1 target gene expression (Supplementary table 5). In the inflamed setting, there was no conforming pattern between the presence or absence of the rs6087990 risk allele and the presence or absence of the NFKB1 or PKCB hubs with regard to NFKB1 target gene expression.

HDAC7 (Histone Deacetylase 7) is an epigenetic regulator, which represses gene expression in muscle maturation by repressing transcription of myocyte enhancer factors such as *MEF2A, MEF2B* and *MEF2C* through deacetylating their histones^48^. The risk allele at rs11168249 is n annotated to regulate HDAC7 both transcriptionally by affecting the TFBS for SMARC4 and post-transcriptionally with a loss of miR-4717 target site on the mRNA of *HDAC7*. In the non-inflamed setting, the presence of the risk allele was associated with significantly higher NFKB1 target gene expression in Clusters 2 and 3 (where NFKB1 was affected by a SNP), with the converse being true in Clusters 1 and 4, where NFKB1 was not affected. This suggests an important relationship between HDAC7 and NFKB1. However, in the inflamed setting, the risk allele at rs11168249 conferred neither protection against, nor significant escalation of NFKB1 target gene expression in any of the clusters.

Overall these observations offer understanding of how non-hub SNP affected proteins may regulate inflammatory response in UC, dependant on patient-specific network footprints represented in the four clusters. Taking into account the SNP profile of patients of these non-hub SNP-affected proteins could lead to further insight when considering therapy, for example targeting LSP1 in patients with PKCB− NFKB1+ network footprints (Cluster 2 patients), but not PKCB− NFKB1− network footprints (Cluster 4 patients), or targeting MAML2 in patients with NFKB1− network footprints (Clusters 1 and 3) but not those with NFKB1+ network footprints (Clusters 2 and 3).

## Discussion

We used UC as a model of a complex genetic disease where there is a need for precision medicine. For this, we designed an integrated systems genomics workflow, termed iSNP, to layer patient data from population wide genomics with network biology and transcriptomics. In doing so we captured the complex genetic background contributing to disease pathogenesis on an individual patient basis.

This study is, to our knowledge, the first documented approach of using functional annotation of non-coding SNPs at an individual patient level. As non-coding SNPs contribute to approximately 90% of disease associated SNPs, analysing them, especially to facilitate precision medicine analysis is of high importance. To do this in a reproducible and automated way, we created a novel, integrative approach (Figure 1) to identify potential functional annotations and stringent quality controls. Quality controls for such computational pipelines are critical as for example, one difficulty with SNP functional analysis is the presence of non-coding SNPs that are tagging SNPs (SNPs with high linkage disequilibrium to other causal SNPs), therefore using them could add false affected proteins (noise) to subsequent network analysis. Although it has been shown that up to 90% of non-coding SNPs are non tagging^26^, to ensure we used the highest quality data in this study we only utilised SNPs which had been fine-mapped either from immunochip or from a publicly available dataset from the Broad Institute^31^.

In terms of SNPs regulating gene expression, we focused on two potential regulatory effects in this study – transcription factor binding sites and miRNA target sites. We acknowledge that other regulatory SNP effects such as splicing sites and SNP effects on long non-coding RNAs are relevant, however for this first study we focused on two regulatory SNP effects that were well grounded in the literature. Using TFBS motif predictions is a common method to annotate SNPs to affect the expression of certain genes, however, in may cases these predictions could contain false positive data. We therefore confirmed that the identified annotations for SNP regulatory effects are consistent with the available literature. In particular, SNP rs608799 is located −283bp from the exon 1A transcription start site in the DNMT3B promoter region and is a CPG rich area^49^, which has been annotated as a transcription factor binding site and prioritised for IBD previously^50^ ; rs11168249 is an intronic SNP (HDAC7) within a known transcription factor binding rich loci, therefore the annotation of the SNP affecting a transcription factor binding site is highly probable. Rs11041476 (affecting LSP1) and rs7404095 (affecting PKCB) are both experimentally validated SNPs affecting miRNA TS^51^.

As an integrative approach, we built upon previous network level studies, which have analysed the cumulative effects of multiple regulatory SNPs^52^. We applied these network approaches for analysis in individual patients, instead of general diseased networks, and tracked the effect of regulatory SNP co-occurrences for each patient. Consequently, we were able to reflect the perturbations of SNPs which otherwise have a low individual effect size^53^, on complex intracellular signaling networks in the individual. To identify the pathogenic effect of these regulatory perturbations, we needed to integrate the SNP annotated genes into a protein-protein interaction network. Using protein-protein interactions and signalling networks to assess pathogenesis is a well-grounded approach. It has been used to identify key disease protein modules^54^ for example in asthma^55^. It has also been applied in many studies to determine hub proteins as the central nodes and drivers of pathogenesis of a disease. Most recently this network approach was used to determine proteins important in asthma disease progression^56^, Parkinson’s disease pathogenesis^57^ and to identify hypertension biomarkers^58^. Beside using interaction networks to identify disease-related modules and key proteins, it has also been used for finding novel pathogenetic players among the interactor partners of already known, key pathogenic genes by “guilt by association”. We previously used this approach to identify potential drug repurposing targets in cancer^28^. In the current study we integrated all these three network reconstruction and analysis methodologies to understand the pathogenesis of complex diseases, such as UC better. A key element in any network biology analysing pipeline is the selection of background interactome network source^59^. To avoid the bias of specific databases, and to maximise the coverage of the networks we are analysing, in this study we used OmniPath^32^. OmniPath integrates information from more than 25 manually curated signalling network resources in a standardized way. Using OmniPath, we also minimised the bias from computational predictions or high-throughput experiments, which may cause inherent ‘noise’ within the networks.

Using the iSNP pipeline for analysing an East Anglian cohort of UC patients, first we focused on identifying UC-related network modules by looking at the protein-protein interaction network affected by the UC SNPs. We identified seven disease associated modules centred around NFKB1, PKCB/FCGR2A, LSP1/GNA12, HDAC7, DNMT3B, and MAML2 (Figure 2). When we analysed the data in a patient-specific way (i.e., reconstructing the networks for each patient separately, we identified PKCB and NFKB1 as two large disease-associated hubs, both of which have been previously associated with UC pathogenesis^60,61^. To identify patient cohorts based on their network footprint, we clustered them by similarity (Figure 3b). For this, we utilised two methods: hierarchical clustering and multidimensional scaling which resulted in the same patient clusters, showing that this outcome was stable with respect to the method employed. This form of unsupervised clustering has been documented^62–64^ and validated in other patient integration network approaches such as NetDx^65^. Despite having good coverage of patient metadata from the UK IBD Genetic Consortium and IBD-Leuven cohorts, supervised clustering did not identify any association with clinical parameters, probably reflecting the relatively low sample sizes in each group. The functional analysis of the patient clusters suggests that different pathogenic pathways are active dependant on patient SNP profiles (Figure 3c). The enriched pathways for the patient clusters also indicate an association between clusters and cell specificity. An example of this is the identification of signalling pathways specific to immune cell types including T cells (Figure 3c, Supplementary Table 3). We propose T cell specificity in Clusters 1 and 3, *via* NFKB1 and indirect involvement through the FC-gamma receptor pathways. This is supported by the literature: NFKB1 is involved in T-cell maturation^66^. FC-gamma receptor related processes, which were an enriched Gene Ontology Biological Process in Clusters 1-3, are also known to affect activation of NFKB1 through NEMO/IKKy^67,68^. Interestingly, we identified a cohort of patients (Cluster 4) whose pathogenesis is driven by non-hub SNP-affected proteins: LSP1, MAML2 and epigenetic modifiers HDAC7 and DNMT3B. These have not been clearly linked with UC pathogenesis previously. We were able to identify potential roles for these proteins in mediating inflammation in patients from genotyped transcriptomic data (Figure 4). Review of the literature gave further insight into how UC pathogenesis may be affected by these SNPs.

LSP1 is intracellular F-actin binding cytoskeletal protein^69^. LSP1 bridges the innate and adaptive immunity, has a role in wound healing, and ingress and intracellular degradation of eukaryotic viruses. We therefore propose that LSP1 acts as a potential interface in UC pathogenesis between the genetic predisposition and environmental signals. In terms of cell specificity, LSP1 is found in mature CD8+ T cells where it reduces the activity of Bim, reducing apoptosis^70^. It is also functions as a negative regulator of cell motility in neutrophils and dendritic cells^71, 72^. Overexpression of LSP1 leads to neutrophil and dendritic cellular rigidity and reduced cell motility. Further work is required to explore the role of LSP1 in the pathogenesis of UC and its potential as a drug target.

MAML2 mediates cross-talk between the inflammatory NFKB1 pathway, and the wound healing Notch pathway. MAML2 is a cofactor in the Notch pathway, facilitating the binding of the active intracellular domain of the NOTCH1 protein to the Notch pathway transcription factor CSL^73^. NOTCH1 itself can also bind to the IKK complex and through it indirectly activating NFKB1^44–46^. We propose that the pathogenic mechanism of the MAML2 SNP via the loss of the miRNA-TS of mir-4495 is to modulate the NOTCH1-NFKB1 cross-talk. This phenomenon was visible in the non-inflamed colonic samples from the Leuven cohort of patients, where we detected significantly decreased NFKB1 target expression (Figure 4).

We have shown that LSP1 and MAML2 affecting SNPs have an impact on downstream NFKB1 target gene expression in the inflamed and non- inflamed colon. These are both targets for further investigation for potential targeted therapy. The two additional non-hub SNP affected proteins HDAC7 and DNMT3B are epigenetic proteins that also showed patient cluster specific changes in NFKB1 target gene expression in the inflamed colon (Figure 4). This further emphasises the known importance of epigenetic regulation in UC^74^. Further work into the role of SNPs affecting epigenetic regulators of the dynamic regulation of pathogenic pathways in UC is required.

The aim of this study was to integrate systems genomics and network biology techniques to bridge the gap between GWAS and individual patients to allow for precision medicine. Whilst due to the available sample size we were not able to identify a link between the individual network footprints and clinical parameters in UC, we have been able to shed further light on UC pathogenesis, and identified new potential targets for precision therapeutics. Peters et al^27^ integrated SNP and RNA variations in IBD without annotating them to identify core immune activation modules. They used Bayesian networks with large IBD cohorts, and identified macrophage cell types as a key player in IBD pathogenesis. We took a different approach in annotating the SNP variations from large cohorts, thereby integrating a functional role to the SNPs with protein-protein networks and signalling networks. We then were able to identify individual patient pathways to disease, which is novel in this field. By broadening the pathogenic pathways from the known immune pathways, we identified pathways which are patient specific and also cell specific, and this is something that will continue to be explored in the future. Integration of multi-omics data and gene networks has been used in schizophrenia, to identify risk genes which enrich in brain tissue for potential drug targeting^75^. Wang et al used stratified linkage disequilibrium to identify risk genes from 100 schizophrenia associated SNPs which they then enriched to brain tissue. Within iSNP we utilised SNPs that had already been enriched to the colon by Fahr et al. Differing from Wang et al, we utilised all the known available SNPs and used protein-protein networks to identify disease associated hubs of proteins, instead of identifying new genes in linkage disequilibrium with SNPs. By doing this, we were able to identify potential novel protein drug targets for specific patient cohorts.

The iSNP workflow is not limited to UC. iSNP is not disease specific and is automated, therefore, can be utilised for analysis of large SNP data repositories. We believe that future, precision medicine works expanding the utility of iSNP into other complex genetic diseases, including Crohn’s disease and other complex, inflammatory diseases such as arthritis, Alzheimer’s disease, autoimmune liver disease and cardiovascular disease is now possible and available for the community.

## Conclusion

We developed the novel integrative SNP Network Platform (iSNP) workflow to identify patient-specific network footprints. These network footprints are based on the regulatory SNP-affected genes and their first neighbour protein-protein interactors. Using iSNP, we have identified how different cellular pathways are associated with UC pathogenesis, and their dependence on the network footprint of individual patients. By combining the iSNP analysis and gene ontology, we determined patient-specific pathways to disease. We identified novel pathways linking the pathogenic effectors of genetic susceptibility, immune modulation, and environmental triggers. Further work into elucidating the exact molecular interactions would allow for patient-specific targeting of these pathogenic pathways. The iSNP workflow has the potential to advance precision medicine by identifying new patient-specific pathogenic pathways and novel personalised drug targets in other complex diseases.

## Methods

### Sources of SNP data

UC associated index SNPs were identified from the UK IBD genetics consortium Immunochip data^9^ and the Broad Institute Repository^31^. If no fine mapping was available for an index SNP (the immunochip finemapped SNP had an R^2^ <0.8) then the highest proxy partners (based on tightest linkage disequilibrium and distance) were assessed using a SNP proxy search and were included in the analysis. Each SNP was annotated using Ensembl from the rsID using the genome map GRCH38.p7. Disease-associated SNPs were retrieved from the original data source.

Using this combined SNP dataset, we compiled UC-specific SNP data for 377 UC patients from seven centres across East Anglia, UK (Cambridge, Norwich, Ipswich, Welwyn Garden City, Luton, Bedford, and West-Suffolk). The examined patients were aged between 25 and 100 years. The mean age of diagnosis was 37 with standard deviation of 14.9 years. 246 patients were on mesalazine treatment and 124 with additional immunomodulatory treatment. For additional data see Supplementary Table 2. SNPs were characterised into different types depending on their location in the genome: exonic (missense, synonymous), intronic/non-translated regions and intergenic. Flanking nucleotide sequences were obtained from dbSNP^76^. For the analysed SNPs see Supplementary Table 1.

### Assessing the effect of SNPs on transcription factor binding sites and miRNA-TS

From the JASPAR database we downloaded 396 human transcription factors’ binding profiles represented by Position Specific Scoring Matrices (PSSMs)^77^. The PSSMs downloaded in JASPAR format were converted to the TRANSFAC format to ease handling of results. To assess the effect of the SNP on the gain or loss of putative TF binding sites, flanking sequences 50 bases upstream and downstream of the SNPs were extracted. The Regulatory Sequence Analysis Tool (RSAT) *matrix-scan*^78^ was used to search for potential TFBS in the ancestral and patient-specific mutant alleles. The background model estimation was determined by using residue probabilities from the input sequences with a Markov order of 1. The search was subject to both strands of the sequences. Hits with a P-value ≤ 1e-05 were considered as binding sites. Other parameters were set at default values.

To assess the effect of the SNPs in miRNA-TSs, the 22bp sequences of mature miRNAs were retrieved from miRBase^79^. The flanking sequences of SNPs were assessed for the presence of miRNA-TSs using miRanda^80^. Hits occurring in the seed region (2’−8’) of the miRNAs and with alignment scores ≥ 90 and energy threshold ≤ −16 kcal/mol were considered as TS. Other parameters were set to default settings. A final manual check was performed to ensure that the SNPs overlapped with the predicted TFBS or miRNA target sites.

We also considered gain or loss of the regulatory interactions between TFs and protein-coding genes in our analysis, where the protein-coding gene was within 10kb upstream or downstream of the SNP-affected TFBS. This information was retrieved using the feature retrieval function of the UCSC genome table browser^81^. We also captured pre-existing regulatory interactions with experimentally determined binding regions/sites. In these cases, the protein coding gene(s) at the cis level corresponding to the SNP were assigned as targets of the TF which recognises the binding regions/sites.

All gains or losses of regulatory interactions and protein coding genes via SNP-affected miRNA-TSs were included in the network except when the SNPs were annotated as intergenic. The effect of SNPs on the uncovered TFBS or miRNA-TSs were classified into either a gain or loss of binding site/target site or a neutral change. Only those sites identified as loss or gain with respect to sites corresponding to the ancestral allele were considered for subsequent analysis. We called the genes corresponding to such SNPs ‘SNP-affected genes’ from here onwards.

### Network construction and analysis

Protein-protein interactions of the proteins encoded by SNP-affected genes were obtained from OmniPath in January 2017^32^. For each patient, the set of proteins encoded by SNP-affected genes and their first interactors (first neighbours) were defined as the UC-associated network footprint of a particular patient. The union of all network footprints, the UC-network, was analysed and visualized in Cytoscape 3.3.0^33^ using the inverted self-organizing map layout. We retained only those SNP-affected genes which were present in the OmniPath resource and, which formed a giant component with their interactors. Patient-specific networks were constructed using the Cytoscape CyRestClient 0.6 in Python^82^.

Cluster analysis was carried out by using the Clustermaker Cytoscape app^83^ implementing the GLay clustering method^84^, which is an implementation of the Girvan-Newman clustering algorithm^85^. Briefly, the clustering method deletes the highest betweenness edges from the network until the network collapsed to non-connected components and these components form the clusters. We call the network clusters from here onwards ‘modules’, to be distinguishable from patient clusters.

### Hierarchical clustering, multidimensional scaling methods and statistical analysis

The *Scipy scikit-learn* package was used for hierarchical clustering^86^ of the patient-specific clusters. The constructed distance matrix between patients was based on the Hamming distance^87^. If a protein was directly or indirectly affected by a SNP, then it was assigned a “1” in a patient. If the protein was not affected, then it was scored as “0”. Multidimensional scaling was conducted in the KNIME environment using the MSA KNIME node^88 89^. We retained only the first three dimensions. The first two dimensions were plotted in Microsoft Excel.

### Gene expression analysis

We used publicly available microarray datasets (GSE73661 and GSE48959), derived from inflamed and non-inflamed colonic biopsies at the IBD centre Leuven, Belgium, in whom Immunochip data were available (Supplementary Table 4). Gene expression was measured on Affymetrix HGU-133 plus2 and Affymetrix Hugene1.0st platforms. The microarray analysis was conducted in R version 3.5.0.. The gene expression data were platform-wise normalised using the robust multi-array average^90^ through the oligo package^91^. Then the probesets were mapped to UniProt IDs from ENSEMBL BioMart using the AnnotationDbi and the biomaRt package^92,93^. The average of gene expression was taken per UniProt ID if multiple probe set was mapped to one specific UniProt ID.

In the case of the inflamed samples there were not enough replicates per platform. To make the two platforms comparable those genes were considered which had probe sets on both the Affymetrix HGU-133 plus2 and on the Hugene1.0st platforms. Subsequently, the mapping to UniProt ID were ranked per sample and rank differences were calculated between classes. The list of NFKB1 target genes were retrieved from the manually curated TRRUST database^94^ (Supplementary Table 7). We then performed a Gene Set Enrichment Analysis^95^. We considered the gene set significant if the GSEA’s Kolmogorov-Smirnov test P-value was below 0.05. All parameters were kept as default.

### Gene Ontology analysis

The Gene Ontology analysis was performed using *pypathway* analysis tool^96^ which implemented the *goatools* package^97^. Each individual patients affected genes were used for enrichment test against the genes in the OmniPath database. The Sidak false discovery calculation was calculated^98^. We considered a Gene Ontology Biological Process term representative for a cluster if it was enriched with corrected q<0.05 significance more than half of the cluster’s patients.

## Supplementary Tables

Supplementary Table 1 List of SNPs in the UKIBD

Supplementary Table 2 Demographics of patients in the East Anglian cohort

Supplementary Table 3 Enriched gene ontologies per clusters

Supplementary Table 4 Demographics of patients in the Leuven cohort

Supplementary Table 5 List of SNPs in the Leuven cohort

Supplementary Table 6 P values of GSEA results

Supplementary Table 7 NFKB GSEA signature from TRRUST database mapped to UniProt IDs

## Declarations

### Availability of data and material

*MTA*

### Author’s contributions

JB, SC and TK designed the iSNP workflow, and wrote the manuscript with DM. JB, DM, PS, MSB, DF, OK and MM developed and automated the workflow. MP provided the East Anglian SNP data and metadata. DM carried out network analysis, and GSEA. KO, AZ and DM were involved in data interpretation. LJH contributed to writing the manuscript. JB, MP, AW and MT provided the clinical insight or clinical data analysis and all contributed to writing the manuscript. AB supervised the work of DM and AZ, and contributed to writing the manuscript. BM and SV provided the gene expression data and contributed to writing the manuscript. All the authors read and approved the final version of the manuscript.

