## Supplementary table 1 for "A systems genomics approach to uncover patient-specific pathogenic pathways and proteins in a complex disease"

**Distribution of patients in the different clusters**

| Cluster | Mean current age (Range) | % Required immunomodulator (n) | % Mesalazine treatment alone (n) | % no treatment data available (n) | % Male (n) | % Female (n) | Mean age at diagnosis (range) |
|---|---|---|---|---|---|---|---|
| Cluster 1 PKCB+, NFKB- | 62.9 (26-99) | 7 (28) | 15 (55) | 0.27 (1) | 10 (37) | 13 (47) | 37.6 (16-81) |
| Cluster 2 PKCB-, NFKB+ | 60.6 (32-90) | 9 (33) | 17 (63) | 0.53 (2) | 16 (59) | 10 (39) | 38.9 (14-77) |
| Cluster 3 PKCB+, NFKB+ | 58.3 (35-87) | 6 (23) | 17 (63) | 0.80 (3) | 13 (47) | 11 (42) | 36.7 (18-75) |
| Cluster 4 PKCB-, NFKB- | 59.5 (25-100) | 11 (40) | 17 (65) | 0.27 (1) | 14 (52) | 14 (52) | 37.7 (9-83) |
| Total patients | 60.3 | 32 (124) | 65 (246) | 1.86 (7) | 52 (195) | 48 (180) | 37.7 |
