## Supplemental table 2 for "A systems genomics approach to uncover patient-specific pathogenic pathways and proteins in a complex disease"

| SNP_ID | SNP type | Name of affected gene +/- 10 kb | Allele difference ? | Influencing TFBS / mirnaBS ? | Is in the giant componenet of the network? |
|---|---|---|---|---|---|
| rs11041476 | intronic | LSP1 | Yes | Yes | Yes |
| rs11168249 | intronic | HDAC7 | Yes | Yes | Yes |
| rs1598859 | intronic | NFKB1 | Yes | Yes | Yes |
| rs1801274 | missense | FCGR2A | Yes | Yes | Yes |
| rs543104 | intronic | MAML2 | Yes | Yes | Yes |
| rs6087990 | UGV | DNMT3B | Yes | Yes | Yes |
| rs7404095 | intronic | PRKCB | Yes | Yes | Yes |
| rs907611 | UGV | LSP1 | Yes | Yes | Yes |
| rs1182188 | intronic | GNA12 | Yes | Yes | Yes |
| rs11229555 | intronic | GLYAT | Yes | Yes | No |
| rs17119 | intronic within ncRNA gene | LOC101928354 | Yes | Yes | No |
| rs254560 | intronic | C5orf66 | Yes | Yes | No |
| rs254562 | intronic | C5orf66 | Yes | Yes | No |
| rs3742130 | synonymous | UBAC2,GPR18 | Yes | Yes | No |
| rs3851228 | intronic within ncRNA gene | LOC107986522,TRAF3IP2-AS1 | Yes | Yes | No |
| rs4243971 | intergenic | -- | Yes | Yes | No |
| rs4743820 | intronic within ncRNA gene | LINC00484,LOC100507103 | Yes | Yes | No |
| rs477515 | intergenic | -- | Yes | Yes | No |
| rs559928 | intergenic | -- | Yes | Yes | No |
| rs913678 | RRV | -- | Yes | Yes | No |
| rs11676348 | intergenic | IL8RB | Yes | No | No |
| rs12568930 | RRV | -- | Yes | No | No |
| rs17085007 | intergenic | -- | Yes | No | No |
| rs4380874 | intergenic | -- | Yes | No | No |
| rs6062504 | intronic | ZGPAT | Yes | No | No |
| rs941823 | intronic | LINC00598 | Yes | No | No |
| rs10228276 | DGV | HOXA13 | No | No | No |
| rs12103 | synonymous | ACAP3,PUSL1,CPSF3L, MIR6727 | No | No | No |
| rs12254167 | intronic | CCNY | No | No | No |
| rs13277237 | intronic | CCDC26 | No | No | No |
| rs17694108 | intergenic | -- | No | No | No |

| | | | | | |
|---|---|---|---|---|---|
| rs17780256 | 3'UTR | SLC39A11, LOC105371887 | No | No | No |
| rs2425019 | intronic | MMP24 | No | No | No |
| rs259964 | intronic | ZNF831 | No | No | No |
| rs267984 | intronic | DAP | No | No | No |
| rs2816958 | intronic | NR5A2 | No | No | No |
| rs3774937 | intronic | NFKB1 | No | No | No |
| rs6940798 | intronic within ncRNA gene | LOC105375070 | No | No | No |
| rs9297145 | intronic | KPNA7 | No | No | No |
| rs943072 | intronic within ncRNA gene | LOC105375070 | No | No | No |

| Source of SNP | Known eQTL |
|---|---|
| BIFM | |
| IBDS ICUC | |
| BIFM | |
| BIFM UCS ICUC | HSPA6 |
| BIFM | |
| ICIBD | DNMT3B |
| IBDS ICIBD | PRKCB |
| IBDS ICIBD BII BIFI | LSP1, TNNI2 |
| ICUC | GNA12 |
| ICIBD | |
| IBDS ICIBD | |
| BIFM BII UCS ICUC | |
| BIFM | |
| ICIBD | |
| IBDS ICIBD | TRAF3IP2 |
| IBDs ICIBD | |
| IBDS ICUC | |
| UCS ICUC | HLA-DRB1,HLA-DQA1,HLA-DQB1 |
| IBDS ICIBD | TRPT1,CCDC88B,FLRT1 |
| IBDS ICIBD | |
| BIFM BII | |
| IBDS ICUC BIFM | |
| IBDS ICUC BIFM BII | |
| ICUC BII BIFM | DLD |
| IBDS ICIBD | LIME1,SLC2A4RG,ZGPAT |
| IBDS ICUC BII BIFM | |
| BIFM | |
| IBDS ICIBD | |
| BIFM | |
| ICIBD | |
| IBDS ICIBD | |

Legend:

| | |
|---|---|
| BIFM | Broad Insitute fine mapped |
| BII | Broad Institute UC index SNP |
| IBDS | GWAS IBD |
| ICIBD | Immunochip IBD |
| ICUC | Immunochip UC |
| UCS | GWAS UC |

| | |
|---|---|
| ICUC BIFM | |
| BIFM | |
| IBDS ICIBD | |
| BIFM | |
| UCS ICUC BII BIFM | |
| ICUC | MANBA |
| BIFM | |
| IBDs ICIBD | SMURF1 |
| BII BIFM | |
