## Supplemental table 3 for "A systems genomics approach to uncover patient-specific pathogenic pathways and proteins in a complex disease"

| GO ID | Name of proces | Count per cluster Sidak correct | | |
| --- | --- | --- | --- | --- |
| | | 1 | 2 | 3 |
| GO:0050853 | regulation of cell communication | 98 | 82 | 89 |
| GO:0090116 | regulation of signaling | 98 | 83 | 89 |
| GO:0006305 | positive regulation of cell communication | 98 | 78 | 89 |
| GO:0006259 | positive regulation of signaling | 98 | 78 | 89 |
| GO:0006306 | regulation of developmental process | 98 | 84 | 82 |
| GO:0032776 | regulation of multicellular organismal process | 98 | 80 | 89 |
| GO:0044728 | regulation of multicellular organismal development | 98 | 83 | 75 |
| GO:0006304 | positive regulation of response to stimulus | 98 | 63 | 89 |
| GO:0006352 | regulation of intracellular signal transduction | 98 | 57 | 89 |
| GO:0002431 | positive regulation of multicellular organismal process | 98 | 68 | 89 |
| GO:0038093 | positive regulation of molecular function | 98 | 82 | 83 |
| GO:0038095 | regulation of signal transduction | 98 | 58 | 89 |
| GO:0038094 | regulation of cell differentiation | 98 | 76 | 80 |
| GO:0038096 | intracellular signal transduction | 98 | 75 | 87 |
| GO:0007186 | activation of immune response | 98 | 53 | 89 |
| GO:0007187 | immune response-activating cell surface receptor signaling pathwa | 98 | 53 | 89 |
| GO:0007249 | immune response-activating signal transduction | 98 | 53 | 89 |
| GO:0000165 | immune response-regulating cell surface receptor signaling pathw | 98 | 53 | 89 |
| GO:0002756 | immune response-regulating signaling pathway | 98 | 53 | 89 |
| GO:0038061 | positive regulation of immune response | 98 | 53 | 89 |
| GO:0061314 | regulation of response to stimulus | 98 | 53 | 89 |
| GO:0007219 | response to chemical | 98 | 84 | 89 |
| GO:0032774 | response to organic substance | 98 | 78 | 89 |
| GO:0016070 | positive regulation of signal transduction | 98 | 54 | 89 |
| GO:0007265 | innate immune response-activating signal transduction | 98 | 53 | 89 |
| GO:0007266 | Fc-epsilon receptor signaling pathway | 98 | 53 | 89 |
| GO:0031295 | regulation of immune system process | 98 | 53 | 89 |
| GO:0050852 | activation of innate immune response | 98 | 51 | 89 |
| GO:0035666 | cellular response to organic substance | 98 | 61 | 89 |
| GO:0000187 | regulation of immune response | 98 | 50 | 89 |
| GO:0002253 | protein phosphorylation | 98 | 70 | 75 |
| GO:0002218 | response to oxygen-containing compound | 98 | 78 | 89 |
| GO:0032147 | antigen receptor-mediated signaling pathway | 98 | 53 | 78 |
| GO:0007188 | signal transduction | 98 | 58 | 89 |
| GO:0048856 | cellular response to chemical stimulus | 98 | 62 | 89 |
| GO:0048513 | positive regulation of intracellular signal transduction | 98 | 41 | 89 |
| GO:0050851 | Fc receptor signaling pathway | 93 | 53 | 74 |
| GO:0006915 | positive regulation of cellular protein metabolic process | 98 | 41 | 89 |
| GO:0019438 | regulation of cellular process | 98 | 66 | 89 |
| GO:0009912 | response to endogenous stimulus | 98 | 73 | 89 |
| GO:0065007 | regulation of cell death | 98 | 64 | 89 |

| | | | | |
|---|---|---|---|---|
| GO:0009058 | stimulatory C-type lectin receptor signaling pathway | 98 | 53 | 89 |
| GO:0048514 | response to nitrogen compound | 98 | 72 | 88 |
| GO:0098751 | innate immune response activating cell surface receptor signaling | 98 | 53 | 89 |
| GO:0055074 | positive regulation of immune system process | 98 | 37 | 89 |
| GO:0055080 | response to organonitrogen compound | 98 | 73 | 79 |
| GO:0001775 | positive regulation of developmental process | 98 | 55 | 80 |
| GO:0008219 | regulation of apoptotic process | 98 | 59 | 89 |
| GO:0030154 | phosphorylation | 98 | 68 | 65 |
| GO:0045165 | cell surface receptor signaling pathway | 98 | 47 | 89 |
| GO:0008283 | positive regulation of cellular process | 98 | 43 | 89 |
| GO:0007166 | positive regulation of protein metabolic process | 98 | 38 | 89 |
| GO:0061311 | regulation of programmed cell death | 98 | 57 | 89 |
| GO:0006725 | positive regulation of protein phosphorylation | 93 | 41 | 77 |
| GO:0044249 | regulation of protein modification process | 97 | 31 | 81 |
| GO:0006874 | positive regulation of I-kappaB kinase/NF-kappaB signaling | 98 | 55 | 89 |
| GO:0030003 | positive regulation of macromolecule metabolic process | 98 | 10 | 89 |
| GO:0055082 | T cell receptor signaling pathway | 98 | 53 | 89 |
| GO:0048869 | regulation of phosphorylation | 98 | 50 | 64 |
| GO:0072503 | regulation of molecular function | 98 | 65 | 73 |
| GO:0019725 | cellular response to oxygen-containing compound | 98 | 45 | 89 |
| GO:0006873 | regulation of cellular protein metabolic process | 98 | 40 | 89 |
| GO:0034645 | regulation of protein phosphorylation | 98 | 44 | 65 |
| GO:0044260 | positive regulation of biological process | 98 | 30 | 89 |
| GO:0044237 | positive regulation of metabolic process | 98 | 13 | 89 |
| GO:0006875 | regulation of I-kappaB kinase/NF-kappaB signaling | 98 | 51 | 89 |
| GO:0044271 | positive regulation of cellular metabolic process | 98 | 13 | 89 |
| GO:0034641 | positive regulation of innate immune response | 98 | 22 | 89 |
| GO:0044267 | positive regulation of nitrogen compound metabolic process | 98 | 12 | 89 |
| GO:0006464 | response to stimulus | 98 | 47 | 89 |
| GO:0071216 | MAPK cascade | 98 | 45 | 89 |
| GO:0070887 | regulation of phosphorus metabolic process | 98 | 48 | 59 |
| GO:0071345 | cellular response to endogenous stimulus | 98 | 41 | 89 |
| GO:0035690 | positive regulation of protein modification process | 91 | 26 | 75 |
| GO:0071495 | macromolecule modification | 87 | 40 | 73 |
| GO:0071496 | regulation of biosynthetic process | 98 | 3 | 89 |
| GO:0071363 | regulation of cellular biosynthetic process | 98 | 3 | 89 |
| GO:0032870 | regulation of gene expression | 98 | 3 | 89 |
| GO:0032869 | regulation of macromolecule biosynthetic process | 98 | 3 | 89 |
| GO:0071396 | positive regulation of phosphorylation | 84 | 36 | 64 |
| GO:1901699 | cellular response to organonitrogen compound | 98 | 41 | 78 |
| GO:0071407 | regulation of cellular macromolecule biosynthetic process | 98 | 3 | 89 |
| GO:0071310 | regulation of RNA biosynthetic process | 98 | | 89 |
| GO:0071417 | regulation of RNA metabolic process | 98 | | 89 |

| | | | | |
|---|---|---|---|---|
| GO:0034599 | regulation of nucleic acid-templated transcription | 98 | | 89 |
| GO:1901701 | regulation of transcription, DNA-templated | 98 | | 89 |
| GO:1901653 | regulation of protein metabolic process | 98 | 26 | 89 |
| GO:0071375 | negative regulation of cell death | 98 | 29 | 89 |
| GO:0034614 | regulation of nucleobase-containing compound metabolic process | 98 | | 89 |
| GO:0051716 | negative regulation of biosynthetic process | 98 | 3 | 89 |
| GO:0033554 | regulation of phosphate metabolic process | 90 | 41 | 59 |
| GO:0035924 | negative regulation of cellular biosynthetic process | 98 | | 89 |
| GO:0048878 | cellular response to stimulus | 98 | 18 | 89 |
| GO:0006325 | positive regulation of cell differentiation | 98 | 34 | 76 |
| GO:0097549 | negative regulation of programmed cell death | 98 | 27 | 89 |
| GO:0034401 | regulation of macromolecule metabolic process | 98 | | 89 |
| GO:0006342 | regulation of nitrogen compound metabolic process | 98 | | 89 |
| GO:0016569 | cellular response to nitrogen compound | 98 | 26 | 89 |
| GO:0019221 | negative regulation of gene expression | 91 | | 89 |
| GO:0002753 | negative regulation of cellular process | 98 | 11 | 89 |
| GO:0006952 | regulation of cellular metabolic process | 98 | | 89 |
| GO:0032502 | regulation of primary metabolic process | 98 | | 89 |
| GO:0072507 | negative regulation of cellular macromolecule biosynthetic proces | 98 | | 89 |
| GO:0042733 | negative regulation of macromolecule biosynthetic process | 98 | | 89 |
| GO:0003160 | regulation of metabolic process | 98 | | 89 |
| GO:0061154 | negative regulation of nitrogen compound metabolic process | 98 | 3 | 89 |
| GO:0007167 | regulation of innate immune response | 98 | 15 | 89 |
| GO:0009913 | negative regulation of apoptotic process | 98 | 21 | 89 |
| GO:0030855 | negative regulation of response to stimulus | 98 | 31 | 78 |
| GO:0050673 | peptidyl-serine modification | 71 | 40 | 71 |
| GO:0061198 | negative regulation of cellular metabolic process | 98 | | 89 |
| GO:0016458 | positive regulation of gene expression | 98 | | 89 |
| GO:0060789 | negative regulation of RNA biosynthetic process | 91 | | 89 |
| GO:0018130 | negative regulation of nucleic acid-templated transcription | 91 | | 89 |
| GO:0046483 | negative regulation of transcription, DNA-templated | 91 | | 89 |
| GO:0070932 | regulation of transcription by RNA polymerase II | 98 | | 89 |
| GO:0070734 | phosphate-containing compound metabolic process | 94 | 58 | 31 |
| GO:0016575 | cellular response to reactive oxygen species | 98 | 18 | 89 |
| GO:0016570 | negative regulation of nucleobase-containing compound metaboli | 91 | | 89 |
| GO:0016572 | I-kappaB kinase/NF-kappaB signaling | 98 | 17 | 89 |
| GO:0042592 | phosphorus metabolic process | 91 | 58 | 31 |
| GO:0002252 | positive regulation of DNA-binding transcription factor activity | 98 | 16 | 89 |
| GO:0002429 | regulation of binding | 98 | 16 | 89 |
| GO:0002757 | negative regulation of RNA metabolic process | 89 | | 89 |
| GO:0002768 | signal transduction by protein phosphorylation | 98 | 18 | 86 |
| GO:0002433 | regulation of DNA-binding transcription factor activity | 98 | 15 | 89 |
| GO:0002764 | positive regulation of defense response | 98 | 10 | 89 |

| | | | | |
|---|---|---|---|---|
| GO:0002376 | negative regulation of biological process | 98 | 2 | 89 |
| GO:0006954 | positive regulation of macromolecule biosynthetic process | 98 | | 89 |
| GO:0002220 | positive regulation of cellular biosynthetic process | 98 | | 89 |
| GO:0002758 | regulation of biological process | 98 | 24 | 75 |
| GO:0060120 | cellular protein modification process | 93 | 29 | 63 |
| GO:0098771 | protein modification process | 93 | 29 | 63 |
| GO:0070498 | positive regulation of RNA biosynthetic process | 98 | | 89 |
| GO:0044419 | positive regulation of nucleic acid-templated transcription | 98 | | 89 |
| GO:0030522 | positive regulation of biosynthetic process | 98 | | 89 |
| GO:0035556 | response to inorganic substance | 86 | 69 | 30 |
| GO:0050801 | positive regulation of transcription by RNA polymerase II | 98 | | 89 |
| GO:0002521 | negative regulation of metabolic process | 83 | | 89 |
| GO:0050900 | negative regulation of macromolecule metabolic process | 80 | | 89 |
| GO:0031294 | positive regulation of RNA metabolic process | 98 | | 89 |
| GO:0009059 | response to stress | 98 | | 89 |
| GO:0098732 | regulation of biological quality | 98 | 82 | 3 |
| GO:0043170 | positive regulation of transcription, DNA-templated | 98 | | 89 |
| GO:0043414 | response to external stimulus | 98 | 20 | 74 |
| GO:0043412 | positive regulation of nucleobase-containing compound metabolic | 98 | | 89 |
| GO:0033598 | cellular response to stress | 98 | | 89 |
| GO:0008152 | response to hormone | 98 | 12 | 78 |
| GO:0055065 | regulation of response to stress | 98 | 1 | 89 |
| GO:0032259 | regulation of cellular component organization | 98 | 84 | |
| GO:0050804 | NIK/NF-kappaB signaling | 98 | | 89 |
| GO:0003159 | apoptotic process | 98 | | 89 |
| GO:0051704 | cell death | 98 | | 89 |
| GO:0030099 | cellular response to hormone stimulus | 98 | | 89 |
| GO:0043392 | cytokine-mediated signaling pathway | 98 | | 89 |
| GO:0043433 | inflammatory response | 98 | | 89 |
| GO:1902679 | interleukin-1-mediated signaling pathway | 98 | | 89 |
| GO:0051253 | mammary gland epithelial cell proliferation | 98 | | 89 |
| GO:0043066 | pattern recognition receptor signaling pathway | 98 | | 89 |
| GO:0048519 | positive regulation of NF-kappaB transcription factor activity | 98 | | 89 |
| GO:0009890 | programmed cell death | 98 | | 89 |
| GO:0010648 | regulation of DNA binding | 98 | | 89 |
| GO:0045786 | regulation of defense response | 98 | | 89 |
| GO:0060548 | regulation of type I interferon production | 98 | | 89 |
| GO:0031327 | response to cytokine | 98 | | 89 |
| GO:2000113 | stress-activated MAPK cascade | 98 | | 89 |
| GO:0031324 | toll-like receptor signaling pathway | 98 | | 89 |
| GO:0048523 | peptidyl-serine phosphorylation | 52 | 41 | 59 |
| GO:0010629 | positive regulation of cell death | 98 | 13 | 71 |
| GO:0045814 | DNA methylation on cytosine | 49 | 40 | 43 |

| | | | | |
|---|---|---|---|---|
| GO:0060965 | peptidyl-amino acid modification | 50 | 21 | 74 |
| GO:0045967 | response to organic cyclic compound | 96 | | 82 |
| GO:1902532 | regulation of cytokine production | 85 | | 89 |
| GO:0010558 | response to abiotic stimulus | 92 | | 84 |
| GO:0010605 | positive regulation of phosphate metabolic process | 73 | 16 | 55 |
| GO:0009892 | positive regulation of phosphorus metabolic process | 73 | 16 | 55 |
| GO:0044092 | response to reactive oxygen species | 86 | 21 | 64 |
| GO:1901215 | negative regulation of cell communication | 98 | 37 | 38 |
| GO:0051172 | immune system process | 78 | 24 | 47 |
| GO:1903507 | response to lipid | 85 | | 87 |
| GO:0045934 | regulation of MAPK cascade | 76 | 5 | 77 |
| GO:0043069 | negative regulation of signaling | 97 | 35 | 37 |
| GO:2001024 | regulation of cellular response to drug | 98 | | 73 |
| GO:0048585 | positive regulation of type I interferon production | 80 | | 89 |
| GO:0009968 | regulation of anatomical structure morphogenesis | 71 | 83 | |
| GO:0023057 | negative regulation of signal transduction | 88 | 21 | 47 |
| GO:0000122 | cell differentiation | 73 | | 83 |
| GO:0045892 | negative regulation of transcription by RNA polymerase II | 51 | | 89 |
| GO:0048663 | positive regulation of catalytic activity | 54 | 45 | 21 |
| GO:0006807 | cellular response to oxidative stress | 85 | | 69 |
| GO:0090304 | platelet activation | 40 | 65 | 13 |
| GO:0097659 | cellular response to organic cyclic compound | 84 | | 67 |
| GO:0034654 | positive regulation of kinase activity | 59 | 28 | 31 |
| GO:0006139 | biological regulation | 97 | 7 | 46 |
| GO:0035872 | positive regulation of transcription of Notch receptor target | 35 | 29 | 45 |
| GO:0070423 | positive regulation of transferase activity | 53 | 33 | 27 |
| GO:1901362 | positive regulation of cytokine production | 53 | | 89 |
| GO:1901360 | regulation of transferase activity | 68 | 38 | 16 |
| GO:1901576 | regulation of DNA-templated transcription in response to stress | 50 | | 89 |
| GO:0071704 | regulation of hemopoiesis | 64 | | 75 |
| GO:1901564 | response to peptide | 98 | 6 | 34 |
| GO:0002221 | rhythmic process | 39 | | 80 |
| GO:0018193 | positive regulation of binding | 98 | | 37 |
| GO:0018209 | cellular macromolecule metabolic process | 44 | | 77 |
| GO:0018105 | response to toxic substance | 49 | 14 | 41 |
| GO:0018210 | cellular response to peptide | 98 | | 31 |
| GO:0018107 | positive regulation of protein kinase activity | 68 | 21 | 21 |
| GO:0038083 | stress-activated protein kinase signaling cascade | 32 | | 89 |
| GO:0018212 | regulation of lipid metabolic process | 64 | 34 | 13 |
| GO:0018108 | positive regulation of cellular component organization | 62 | 30 | 20 |
| GO:0006796 | cellular protein metabolic process | 65 | | 48 |
| GO:0006793 | regulation of cellular response to stress | 22 | | 89 |
| GO:0016310 | response to drug | 28 | | 70 |

| | | | | |
|---|---|---|---|---|
| GO:0030168 | developmental process | 64 | 45 | |
| GO:0051091 | regulation of kinase activity | 47 | 36 | 9 |
| GO:0043123 | regulation of transcription from RNA polymerase II promoter in re: | 18 | | 88 |
| GO:0043406 | regulation of protein kinase activity | 64 | 26 | 10 |
| GO:0043410 | response to growth factor | 56 | | 45 |
| GO:0051092 | macromolecule metabolic process | 16 | | 68 |
| GO:0045747 | positive regulation of cell population proliferation | 19 | | 64 |
| GO:1902680 | protein metabolic process | 39 | | 56 |
| GO:0051254 | chromatin organization | 8 | | 67 |
| GO:0046579 | intracellular receptor signaling pathway | 13 | | 87 |
| GO:0035025 | peptidyl-threonine phosphorylation | 39 | 41 | 18 |
| GO:0050870 | defense response | 52 | | 46 |
| GO:0045766 | T cell costimulation | 37 | 53 | |
| GO:0043065 | lymphocyte costimulation | 37 | 53 | |
| GO:0051099 | peptidyl-threonine modification | 39 | 41 | 13 |
| GO:0048518 | interspecies interaction between organisms | 42 | | 54 |
| GO:0009891 | regulation of oxidative stress-induced cell death | 80 | | 18 |
| GO:0043536 | regulation of response to drug | 98 | | |
| GO:0043085 | regulation of catalytic activity | 50 | 20 | 14 |
| GO:0050867 | regulation of postsynaptic membrane potential | | 83 | |
| GO:0010647 | covalent chromatin modification | 2 | | 63 |
| GO:0010942 | response to oxidative stress | 41 | | 50 |
| GO:0010720 | RNA biosynthetic process | 5 | | 79 |
| GO:0045597 | DNA alkylation | | 2 | 43 |
| GO:0008284 | DNA methylation | | 2 | 43 |
| GO:0031328 | cellular calcium ion homeostasis | 43 | 38 | |
| GO:0051130 | calcium ion homeostasis | 43 | 38 | |
| GO:0031325 | cellular response to growth factor stimulus | 42 | 9 | 29 |
| GO:0048522 | positive regulation of apoptotic process | 59 | 2 | 28 |
| GO:1903829 | positive regulation of programmed cell death | 59 | 2 | 28 |
| GO:0032270 | cellular divalent inorganic cation homeostasis | 43 | 37 | |
| GO:1905269 | regulation of lymphocyte activation | 52 | | 35 |
| GO:2001252 | regulation of cytosolic calcium ion concentration | 43 | 30 | |
| GO:0001819 | positive regulation of MAPK cascade | 32 | | 50 |
| GO:0007204 | histone modification | | | 63 |
| GO:0051482 | response to biotic stimulus | 10 | | 72 |
| GO:0031349 | divalent inorganic cation homeostasis | 43 | 34 | |
| GO:0051094 | regulation of neuron death | 88 | | |
| GO:0010628 | regulation of cellular localization | 32 | 48 | |
| GO:0014015 | DNA methylation or demethylation | | | 39 |
| GO:0031058 | response to metal ion | 39 | 41 | |
| GO:0051345 | negative regulation of gene expression, epigenetic | | | 38 |
| GO:0050778 | regulation of gene expression, epigenetic | | | 43 |

| | | | | |
|---|---|---|---|---|
| GO:0002684 | cellular response to peptide hormone stimulus | 52 | | 31 |
| GO:0045089 | regulation of viral transcription | | | 78 |
| GO:1902533 | regulation of T cell activation | 46 | | 34 |
| GO:0033674 | protein autophosphorylation | 11 | 27 | 9 |
| GO:0002696 | cellular metal ion homeostasis | 43 | 30 | |
| GO:1903039 | regulation of localization | 31 | 43 | |
| GO:1902107 | positive regulation of protein serine/threonine kinase activity | 37 | 12 | 19 |
| GO:0060193 | positive regulation of cytosolic calcium ion concentration | 35 | 29 | |
| GO:0051251 | regulation of cellular protein localization | 36 | 40 | |
| GO:0010557 | cellular cation homeostasis | 43 | 30 | |
| GO:0010604 | regulation of myeloid cell differentiation | 8 | | 68 |
| GO:0009893 | response to UV | 34 | | 44 |
| GO:0044093 | multi-organism process | 29 | | 48 |
| GO:0051240 | cellular ion homeostasis | 41 | 30 | |
| GO:0048636 | metal ion homeostasis | 41 | 30 | |
| GO:1901863 | regulation of vasculature development | 47 | 12 | 18 |
| GO:0051962 | positive regulation of MAP kinase activity | 46 | 10 | 15 |
| GO:0050769 | nitrogen compound metabolic process | 9 | | 55 |
| GO:1901216 | cellular chemical homeostasis | 43 | 30 | |
| GO:0010976 | ion homeostasis | 43 | 30 | |
| GO:0051173 | G protein-coupled receptor signaling pathway | 7 | 30 | |
| GO:1903508 | primary metabolic process | 7 | | 51 |
| GO:0045935 | symbiont process | 27 | | 41 |
| GO:0045937 | viral process | 27 | | 41 |
| GO:0010562 | activation of protein kinase activity | 49 | 1 | 23 |
| GO:0042327 | enzyme linked receptor protein signaling pathway | 13 | 27 | |
| GO:0043068 | chemical homeostasis | 39 | 30 | |
| GO:0045860 | positive regulation of vasculature development | 60 | 12 | |
| GO:0051247 | cation homeostasis | 35 | 29 | |
| GO:0031401 | regulation of transport | 40 | 28 | |
| GO:0001934 | response to external biotic stimulus | 9 | | 58 |
| GO:0071902 | regulation of cell shape | 9 | 54 | |
| GO:0045862 | organic substance metabolic process | 7 | | 49 |
| GO:0048584 | macromolecule biosynthetic process | 1 | | 63 |
| GO:0009967 | homeostatic process | 35 | 31 | |
| GO:0023056 | regulation of cell morphogenesis | | 60 | |
| GO:0051057 | nucleic acid metabolic process | | | 61 |
| GO:0045844 | regulation of MAP kinase activity | 31 | 13 | 13 |
| GO:0045944 | regulation of cell development | 17 | 41 | |
| GO:1901522 | response to lipopolysaccharide | 5 | | 58 |
| GO:0007221 | inorganic ion homeostasis | 30 | 29 | |
| GO:0045893 | nucleobase-containing compound biosynthetic process | | | 58 |
| GO:0051347 | cellular response to drug | 15 | | 40 |

| | | | | |
|---|---|---|---|---|
| GO:0051050 | nucleobase-containing compound metabolic process | | | 57 |
| GO:0032481 | regulation of protein localization | 22 | 28 | |
| GO:1904018 | cellular developmental process | 39 | | 22 |
| GO:0050434 | response to antibiotic | 10 | 17 | 15 |
| GO:0044238 | regulation of DNA metabolic process | | | 57 |
| GO:0012501 | negative regulation of intracellular signal transduction | 37 | 12 | 8 |
| GO:0046777 | regulation of peptidyl-tyrosine phosphorylation | | 52 | |
| GO:0006476 | C-5 methylation of cytosine | | | 14 |
| GO:0035601 | regulation of Notch signaling pathway | 5 | | 29 |
| GO:0019538 | cellular homeostasis | 27 | 27 | |
| GO:0036211 | peptidyl-tyrosine autophosphorylation | | 2 | 9 |
| GO:0006468 | nucleic acid-templated transcription | | | 53 |
| GO:0006508 | transcription, DNA-templated | | | 53 |
| GO:0051101 | positive regulation of leukocyte activation | 52 | | 5 |
| GO:0051052 | response to insulin | 52 | | 4 |
| GO:0051090 | response to peptide hormone | 47 | | 8 |
| GO:0043620 | response to molecule of bacterial origin | 2 | | 49 |
| GO:0043122 | positive regulation of proteolysis | 29 | | 24 |
| GO:0043405 | negative regulation of molecular function | 55 | | |
| GO:0043408 | positive regulation of cell activation | 52 | | 2 |
| GO:1901222 | regulation of protein serine/threonine kinase activity | 26 | 8 | 9 |
| GO:0008593 | positive regulation of lymphocyte activation | 47 | | 5 |
| GO:2001141 | response to mechanical stimulus | 52 | | |
| GO:0051252 | transmembrane receptor protein tyrosine kinase signaling pathway | | 30 | |
| GO:0046578 | positive regulation of neurogenesis | 26 | 10 | 6 |
| GO:0035023 | negative regulation of response to drug | 51 | | |
| GO:0050863 | Fc receptor mediated stimulatory signaling pathway | | | 1 |
| GO:0022603 | Fc-gamma receptor signaling pathway | | | 1 |
| GO:0045765 | Fc-gamma receptor signaling pathway involved in phagocytosis | | | 1 |
| GO:0042981 | immune response-regulating cell surface receptor signaling pathway involved in phagoc | | | 1 |
| GO:0051098 | positive regulation of Notch signaling pathway | | | 24 |
| GO:0050789 | metabolic process | 1 | | 40 |
| GO:0065008 | heterocycle biosynthetic process | | | 41 |
| GO:0009889 | DNA modification | | | 4 |
| GO:0030193 | regulation of cellular response to oxidative stress | 37 | | 8 |
| GO:0043535 | positive regulation of cellular protein localization | 21 | 21 | |
| GO:0050878 | heterocycle metabolic process | | | 39 |
| GO:0051924 | cell activation | 13 | 13 | 2 |
| GO:0050790 | regulation of gliogenesis | | | 25 |
| GO:0050865 | aromatic compound biosynthetic process | | | 38 |
| GO:0010646 | cellular aromatic compound metabolic process | | | 38 |
| GO:0071156 | DNA-templated transcription, initiation | | | 21 |
| GO:0010941 | nucleotide-binding domain, leucine rich repeat containing receptor signaling pathway | | | 39 |

| | | | | |
|---|---|---|---|---|
| GO:0060284 | nucleotide-binding oligomerization domain containing signaling pathway | | | 39 |
| GO:0045595 | regulation of cell population proliferation | | | 39 |
| GO:0022604 | positive regulation of chromatin organization | | | 22 |
| GO:0042127 | positive regulation of T cell activation | 40 | | |
| GO:0031344 | fungiform papilla formation | | | 14 |
| GO:2000136 | organic cyclic compound biosynthetic process | | | 34 |
| GO:0008360 | positive regulation of nervous system development | 21 | 10 | 1 |
| GO:0034248 | positive regulation of cell development | 13 | 8 | 6 |
| GO:0031326 | Notch signaling pathway | | | 17 |
| GO:0044087 | regulation of angiogenesis | 26 | 1 | 9 |
| GO:0051128 | regulation of small GTPase mediated signal transduction | | 3 | |
| GO:0060341 | epithelial cell differentiation | 17 | | 18 |
| GO:2000112 | regulation of blood vessel endothelial cell migration | 16 | 5 | 12 |
| GO:0031323 | response to radiation | 14 | | 20 |
| GO:0050794 | cellular nitrogen compound biosynthetic process | | | 31 |
| GO:1903827 | response to hydrogen peroxide | 5 | | 20 |
| GO:0032268 | regulation of leukocyte activation | 34 | | 1 |
| GO:2001038 | regulation of cell activation | 27 | | 6 |
| GO:0090287 | regulation of response to oxidative stress | 28 | | 5 |
| GO:1900034 | cellular nitrogen compound metabolic process | | | 29 |
| GO:1900407 | transcription initiation from RNA polymerase II promoter | | | 11 |
| GO:0080135 | response to light stimulus | 30 | | 2 |
| GO:1902275 | response to alcohol | 10 | 7 | 6 |
| GO:0050818 | positive regulation of hydrolase activity | 15 | 12 | |
| GO:0001817 | organic cyclic compound metabolic process | | | 27 |
| GO:0050707 | chromatin organization involved in regulation of transcription | | | 21 |
| GO:0001959 | regulation of nervous system development | 5 | 22 | |
| GO:0051480 | negative regulation of neuron death | 29 | | |
| GO:0031347 | cellular response to lipid | 17 | | 11 |
| GO:0050793 | organonitrogen compound metabolic process | 7 | | 20 |
| GO:0070201 | organic substance biosynthetic process | | | 26 |
| GO:0010468 | regulation of inflammatory response | 22 | | 4 |
| GO:0006349 | activation of MAPK activity | 12 | | 12 |
| GO:0040029 | biosynthetic process | | | 23 |
| GO:0014013 | regulation of symbiosis, encompassing mutualism through parasitism | | | 22 |
| GO:1903706 | cellular metabolic process | | | 20 |
| GO:1900046 | positive regulation of leukocyte differentiation | | | 21 |
| GO:0031056 | regulation of Ras protein signal transduction | | 4 | |
| GO:0050776 | cell population proliferation | 22 | | |
| GO:0002682 | positive regulation of angiogenesis | 22 | | |
| GO:0050727 | cellular macromolecule biosynthetic process | | | 20 |
| GO:0045088 | positive regulation of histone modification | | | 14 |
| GO:1902531 | hair follicle placode formation | | | |

| | | | | |
|---|---|---|---|---|
| GO:0033143 | macromolecule methylation | | | |
| GO:0043549 | positive regulation of Rho protein signal transduction | | | |
| GO:0002694 | regulation of tumor necrosis factor-mediated signaling pathway | 21 | | |
| GO:1902105 | regulation of histone modification | | | 13 |
| GO:0060191 | positive regulation of chromosome organization | | | 7 |
| GO:0019216 | methylation | | | |
| GO:0032879 | cellular biosynthetic process | | | 18 |
| GO:0051249 | anatomical structure development | | 16 | |
| GO:0010556 | regulation of neurogenesis | | 16 | |
| GO:0060255 | cellular response to external stimulus | 11 | | 7 |
| GO:0019222 | regulation of leukocyte differentiation | | | 17 |
| GO:1904526 | cellular response to cytokine stimulus | 9 | | 8 |
| GO:0065009 | TRIF-dependent toll-like receptor signaling pathway | | | 16 |
| GO:0043900 | regulation of viral process | | | 16 |
| GO:2000026 | response to tumor necrosis factor | | | 16 |
| GO:0051239 | positive regulation of neuron death | 2 | | 14 |
| GO:0045637 | regulation of response to external stimulus | 15 | | 2 |
| GO:0002761 | tube morphogenesis | | 4 | |
| GO:0051960 | RNA metabolic process | | | 14 |
| GO:0050767 | cytoplasmic pattern recognition receptor signaling pathway | | | 14 |
| GO:1901214 | MyD88-independent toll-like receptor signaling pathway | | | 13 |
| GO:0045664 | leukocyte differentiation | | | 13 |
| GO:0010975 | cellular response to insulin stimulus | 14 | | |
| GO:0051171 | G protein-coupled receptor signaling pathway, coupled to cyclic nucleotide second messenger | | | |
| GO:1903506 | positive regulation of cytosolic calcium ion concentration involved in phospholipase C-activating G | | | |
| GO:0019219 | positive regulation of small GTPase mediated signal transduction | | | |
| GO:0030278 | positive regulation of viral transcription | | | 12 |
| GO:0045667 | response to muscle stretch | | | 12 |
| GO:1903201 | positive regulation of Ras protein signal transduction | | | |
| GO:0090087 | modulation of chemical synaptic transmission | | 10 | |
| GO:0050730 | regulation of trans-synaptic signaling | | 10 | |
| GO:0019220 | Notch signaling involved in heart development | | | |
| GO:0010517 | cell fate commitment | | | |
| GO:0051174 | peptidyl-tyrosine modification | | | |
| GO:0042325 | peptidyl-tyrosine phosphorylation | | | |
| GO:0120035 | vascular endothelial growth factor receptor signaling pathway | | | |
| GO:0060078 | bone cell development | 11 | | |
| GO:1902893 | positive regulation of gliogenesis | | | 10 |
| GO:0080090 | regulation of plasma membrane bounded cell projection organization | | 9 | |
| GO:0043067 | myeloid cell differentiation | | | 9 |
| GO:0043393 | regulation of cell projection organization | | 8 | |
| GO:0045859 | B cell receptor signaling pathway | | | |
| GO:0032880 | regulation of cytokine-mediated signaling pathway | 9 | | |

| | | | | |
|---|---|---|---|---|
| GO:1900180 | response to ethanol | | 6 | |
| GO:0051246 | DNA metabolic process | | | 8 |
| GO:0031399 | epithelial cell proliferation | | | 8 |
| GO:0001932 | regulation of intracellular steroid hormone receptor signaling pathway | | | 8 |
| GO:0071900 | regulation of myeloid leukocyte differentiation | | | 8 |
| GO:0051223 | adenylate cyclase-modulating G protein-coupled receptor signaling pathway | | | |
| GO:0061097 | cell surface receptor signaling pathway involved in heart development | | | |
| GO:2001023 | regulation of Rho protein signal transduction | | | |
| GO:0032101 | regulation of lipase activity | | 7 | |
| GO:1902882 | regulation of phospholipase activity | | 7 | |
| GO:0048583 | regulation of blood coagulation | | 1 | |
| GO:0080134 | regulation of hemostasis | | 1 | |
| GO:0009966 | regulation of multi-organism process | | | 7 |
| GO:0023051 | Ras protein signal transduction | | | |
| GO:0051056 | epidermal cell differentiation | | | |
| GO:0043903 | gene silencing | | | |
| GO:0044057 | immune effector process | | | |
| GO:0099177 | regulation of system process | | 6 | |
| GO:0006357 | endothelial tube morphogenesis | | | |
| GO:0043618 | histone H3-K27 methylation | | | |
| GO:0006355 | leukocyte migration | | | |
| GO:0051338 | morphogenesis of an endothelium | | | |
| GO:0051049 | regulation of establishment of protein localization | | | |
| GO:0010803 | positive regulation of leukocyte cell-cell adhesion | 6 | | |
| GO:0032479 | positive regulation of neuron projection development | | 5 | |
| GO:1901342 | regulation of cellular response to growth factor stimulus | | 5 | |
| GO:0050792 | regulation of neuron projection development | | 5 | |
| GO:0046782 | Rho protein signal transduction | | | |
| GO:0009411 | histone H3 deacetylation | | | |
| GO:0009628 | positive regulation of lipase activity | | | |
| GO:0001101 | positive regulation of muscle organ development | | | |
| GO:0097305 | positive regulation of muscle tissue development | | | |
| GO:0046677 | positive regulation of striated muscle tissue development | | | |
| GO:0009607 | regulation of cell proliferation involved in heart morphogenesis | | | |
| GO:0042221 | regulation of chromatin organization | | | |
| GO:0034097 | proteolysis | | | 5 |
| GO:0042493 | regulation of cell cycle arrest | | | 5 |
| GO:0009719 | positive regulation of blood vessel endothelial cell migration | 5 | | |
| GO:0045471 | cellular response to vascular endothelial growth factor stimulus | 4 | | |
| GO:0043207 | regulation of protein tyrosine kinase activity | | 4 | |
| GO:0009605 | regulation of coagulation | | | |
| GO:0070848 | cellular response to biotic stimulus | | | 4 |
| GO:0009725 | transcription by RNA polymerase II | | | 4 |

| | | | | |
|---|---|---|---|---|
| GO:0042542 | histone deacetylation | | | |
| GO:0010035 | macromolecule deacylation | | | |
| GO:0032868 | protein deacetylation | | | |
| GO:0009416 | protein deacylation | | | |
| GO:0033993 | regulation of peptide transport | | | |
| GO:0032496 | regulation of protein transport | | | |
| GO:0009612 | regulation of neuron differentiation | | 3 | |
| GO:0010038 | negative regulation of DNA-binding transcription factor activity | | | 3 |
| GO:0002237 | positive regulation of transcription from RNA polymerase II promoter involved in cellula | | | 3 |
| GO:0035994 | regulation of NIK/NF-kappaB signaling | | | 3 |
| GO:1901698 | regulation of cellular response to heat | | | 3 |
| GO:0014070 | regulation of pri-miRNA transcription by RNA polymerase II | | | 3 |
| GO:0010033 | positive regulation of transport | 3 | | |
| GO:0010243 | chromatin organization involved in negative regulation of transcription | | | |
| GO:0006979 | chromatin silencing | | | |
| GO:1901700 | embryonic digit morphogenesis | | | |
| GO:1901652 | regulation of body fluid levels | | | |
| GO:0043434 | regulation of microtubule binding | | | |
| GO:0009314 | regulation of cellular amide metabolic process | | 2 | |
| GO:0000302 | histone phosphorylation | | | 2 |
| GO:0050896 | negative regulation of gene silencing by miRNA | | | 2 |
| GO:0006950 | animal organ development | 2 | | |
| GO:0009636 | negative regulation of DNA binding | 2 | | |
| GO:0034612 | regulation of protein binding | 2 | | |
| GO:0048511 | regulation of gene expression by genetic imprinting | | | |
| GO:0046903 | regulation of ossification | | | |
| GO:0032940 | regulation of osteoblast differentiation | | | |
| GO:0007165 | regulation of protein localization to nucleus | | | |
| GO:0023014 | regulation of calcium ion transport | | 1 | |
| GO:0002223 | secretion | | 1 | |
| GO:0051403 | secretion by cell | | 1 | |
| GO:0031098 | negative regulation of cell cycle | | | 1 |
| GO:0044403 | response to acid chemical | | | 1 |
| GO:0002224 | regulation of cellular component biogenesis | 1 | | |
| GO:0006366 | auditory receptor cell fate commitment | | | |
| GO:0006367 | blood vessel morphogenesis | | | |
| GO:0006351 | endocardium morphogenesis | | | |
| GO:0007169 | inner ear receptor cell fate commitment | | | |
| GO:0035239 | negative regulation of growth rate | | | |
| GO:0048010 | neuron fate commitment | | | |
| GO:0016032 | regulation of cytokine secretion | | | |

| ed p<0.05 | Percentage of patinets per clusters | | | | Is the given GO cluster | | | |
|---|---|---|---|---|---|---|---|---|
| 4 | 1 | 2 | 3 | 4 | 1 | 2 | 3 | 4 |
| 42 | 100.00% | 97.62% | 98.89% | 39.62% | 1 | 1 | 1 | 0 |
| 40 | 100.00% | 98.81% | 98.89% | 37.74% | 1 | 1 | 1 | 0 |
| 45 | 100.00% | 92.86% | 98.89% | 42.45% | 1 | 1 | 1 | 0 |
| 45 | 100.00% | 92.86% | 98.89% | 42.45% | 1 | 1 | 1 | 0 |
| 45 | 100.00% | 100.00% | 91.11% | 42.45% | 1 | 1 | 1 | 0 |
| 38 | 100.00% | 95.24% | 98.89% | 35.85% | 1 | 1 | 1 | 0 |
| 49 | 100.00% | 98.81% | 83.33% | 46.23% | 1 | 1 | 1 | 0 |
| 54 | 100.00% | 75.00% | 98.89% | 50.94% | 1 | 1 | 1 | 1 |
| 58 | 100.00% | 67.86% | 98.89% | 54.72% | 1 | 1 | 1 | 1 |
| 42 | 100.00% | 80.95% | 98.89% | 39.62% | 1 | 1 | 1 | 0 |
| 31 | 100.00% | 97.62% | 92.22% | 29.25% | 1 | 1 | 1 | 0 |
| 51 | 100.00% | 69.05% | 98.89% | 48.11% | 1 | 1 | 1 | 0 |
| 36 | 100.00% | 90.48% | 88.89% | 33.96% | 1 | 1 | 1 | 0 |
| 29 | 100.00% | 89.29% | 96.67% | 27.36% | 1 | 1 | 1 | 0 |
| 53 | 100.00% | 63.10% | 98.89% | 50.00% | 1 | 1 | 1 | 0 |
| 53 | 100.00% | 63.10% | 98.89% | 50.00% | 1 | 1 | 1 | 0 |
| 53 | 100.00% | 63.10% | 98.89% | 50.00% | 1 | 1 | 1 | 0 |
| 53 | 100.00% | 63.10% | 98.89% | 50.00% | 1 | 1 | 1 | 0 |
| 53 | 100.00% | 63.10% | 98.89% | 50.00% | 1 | 1 | 1 | 0 |
| 53 | 100.00% | 63.10% | 98.89% | 50.00% | 1 | 1 | 1 | 0 |
| 53 | 100.00% | 63.10% | 98.89% | 50.00% | 1 | 1 | 1 | 0 |
| 12 | 100.00% | 100.00% | 98.89% | 11.32% | 1 | 1 | 1 | 0 |
| 17 | 100.00% | 92.86% | 98.89% | 16.04% | 1 | 1 | 1 | 0 |
| 47 | 100.00% | 64.29% | 98.89% | 44.34% | 1 | 1 | 1 | 0 |
| 48 | 100.00% | 63.10% | 98.89% | 45.28% | 1 | 1 | 1 | 0 |
| 46 | 100.00% | 63.10% | 98.89% | 43.40% | 1 | 1 | 1 | 0 |
| 45 | 100.00% | 63.10% | 98.89% | 42.45% | 1 | 1 | 1 | 0 |
| 44 | 100.00% | 60.71% | 98.89% | 41.51% | 1 | 1 | 1 | 0 |
| 31 | 100.00% | 72.62% | 98.89% | 29.25% | 1 | 1 | 1 | 0 |
| 44 | 100.00% | 59.52% | 98.89% | 41.51% | 1 | 1 | 1 | 0 |
| 33 | 100.00% | 83.33% | 83.33% | 31.13% | 1 | 1 | 1 | 0 |
| 5 | 100.00% | 92.86% | 98.89% | 4.72% | 1 | 1 | 1 | 0 |
| 49 | 100.00% | 63.10% | 86.67% | 46.23% | 1 | 1 | 1 | 0 |
| 29 | 100.00% | 69.05% | 98.89% | 27.36% | 1 | 1 | 1 | 0 |
| 21 | 100.00% | 73.81% | 98.89% | 19.81% | 1 | 1 | 1 | 0 |
| 46 | 100.00% | 48.81% | 98.89% | 43.40% | 1 | 0 | 1 | 0 |
| 53 | 94.90% | 63.10% | 82.22% | 50.00% | 1 | 1 | 1 | 0 |
| 45 | 100.00% | 48.81% | 98.89% | 42.45% | 1 | 0 | 1 | 0 |
| 10 | 100.00% | 78.57% | 98.89% | 9.43% | 1 | 1 | 1 | 0 |
| 1 | 100.00% | 86.90% | 98.89% | 0.94% | 1 | 1 | 1 | 0 |
| 12 | 100.00% | 76.19% | 98.89% | 11.32% | 1 | 1 | 1 | 0 |

| | | | | | | | | |
|---|---|---|---|---|---|---|---|---|
| 25 | 100.00% | 63.10% | 98.89% | 23.58% | 1 | 1 | 1 | 0 |
| 2 | 100.00% | 85.71% | 97.78% | 1.89% | 1 | 1 | 1 | 0 |
| 24 | 100.00% | 63.10% | 98.89% | 22.64% | 1 | 1 | 1 | 0 |
| 43 | 100.00% | 44.05% | 98.89% | 40.57% | 1 | 0 | 1 | 0 |
| 7 | 100.00% | 86.90% | 87.78% | 6.60% | 1 | 1 | 1 | 0 |
| 26 | 100.00% | 65.48% | 88.89% | 24.53% | 1 | 1 | 1 | 0 |
| 10 | 100.00% | 70.24% | 98.89% | 9.43% | 1 | 1 | 1 | 0 |
| 26 | 100.00% | 80.95% | 72.22% | 24.53% | 1 | 1 | 1 | 0 |
| 23 | 100.00% | 55.95% | 98.89% | 21.70% | 1 | 1 | 1 | 0 |
| 28 | 100.00% | 51.19% | 98.89% | 26.42% | 1 | 1 | 1 | 0 |
| 34 | 100.00% | 45.24% | 98.89% | 32.08% | 1 | 0 | 1 | 0 |
| 9 | 100.00% | 67.86% | 98.89% | 8.49% | 1 | 1 | 1 | 0 |
| 46 | 94.90% | 48.81% | 85.56% | 43.40% | 1 | 0 | 1 | 0 |
| 44 | 98.98% | 36.90% | 90.00% | 41.51% | 1 | 0 | 1 | 0 |
| 3 | 100.00% | 65.48% | 98.89% | 2.83% | 1 | 1 | 1 | 0 |
| 58 | 100.00% | 11.90% | 98.89% | 54.72% | 1 | 0 | 1 | 1 |
| 2 | 100.00% | 63.10% | 98.89% | 1.89% | 1 | 1 | 1 | 0 |
| 35 | 100.00% | 59.52% | 71.11% | 33.02% | 1 | 1 | 1 | 0 |
| 4 | 100.00% | 77.38% | 81.11% | 3.77% | 1 | 1 | 1 | 0 |
| 10 | 100.00% | 53.57% | 98.89% | 9.43% | 1 | 1 | 1 | 0 |
| 16 | 100.00% | 47.62% | 98.89% | 15.09% | 1 | 0 | 1 | 0 |
| 39 | 100.00% | 52.38% | 72.22% | 36.79% | 1 | 1 | 1 | 0 |
| 28 | 100.00% | 35.71% | 98.89% | 26.42% | 1 | 0 | 1 | 0 |
| 49 | 100.00% | 15.48% | 98.89% | 46.23% | 1 | 0 | 1 | 0 |
| | 100.00% | 60.71% | 98.89% | 0.00% | 1 | 1 | 1 | 0 |
| 46 | 100.00% | 15.48% | 98.89% | 43.40% | 1 | 0 | 1 | 0 |
| 34 | 100.00% | 26.19% | 98.89% | 32.08% | 1 | 0 | 1 | 0 |
| 45 | 100.00% | 14.29% | 98.89% | 42.45% | 1 | 0 | 1 | 0 |
| | 100.00% | 55.95% | 98.89% | 0.00% | 1 | 1 | 1 | 0 |
| | 100.00% | 53.57% | 98.89% | 0.00% | 1 | 1 | 1 | 0 |
| 29 | 100.00% | 57.14% | 65.56% | 27.36% | 1 | 1 | 1 | 0 |
| | 100.00% | 48.81% | 98.89% | 0.00% | 1 | 0 | 1 | 0 |
| 41 | 92.86% | 30.95% | 83.33% | 38.68% | 1 | 0 | 1 | 0 |
| 30 | 88.78% | 47.62% | 81.11% | 28.30% | 1 | 0 | 1 | 0 |
| 39 | 100.00% | 3.57% | 98.89% | 36.79% | 1 | 0 | 1 | 0 |
| 39 | 100.00% | 3.57% | 98.89% | 36.79% | 1 | 0 | 1 | 0 |
| 39 | 100.00% | 3.57% | 98.89% | 36.79% | 1 | 0 | 1 | 0 |
| 39 | 100.00% | 3.57% | 98.89% | 36.79% | 1 | 0 | 1 | 0 |
| 41 | 85.71% | 42.86% | 71.11% | 38.68% | 1 | 0 | 1 | 0 |
| 2 | 100.00% | 48.81% | 86.67% | 1.89% | 1 | 0 | 1 | 0 |
| 36 | 100.00% | 3.57% | 98.89% | 33.96% | 1 | 0 | 1 | 0 |
| 39 | 100.00% | 0.00% | 98.89% | 36.79% | 1 | 0 | 1 | 0 |
| 39 | 100.00% | 0.00% | 98.89% | 36.79% | 1 | 0 | 1 | 0 |

| | | | | | | | | |
|---|---|---|---|---|---|---|---|---|
| 39 | 100.00% | 0.00% | 98.89% | 36.79% | 1 | 0 | 1 | 0 |
| 39 | 100.00% | 0.00% | 98.89% | 36.79% | 1 | 0 | 1 | 0 |
| 6 | 100.00% | 30.95% | 98.89% | 5.66% | 1 | 0 | 1 | 0 |
| 1 | 100.00% | 34.52% | 98.89% | 0.94% | 1 | 0 | 1 | 0 |
| 37 | 100.00% | 0.00% | 98.89% | 34.91% | 1 | 0 | 1 | 0 |
| 33 | 100.00% | 3.57% | 98.89% | 31.13% | 1 | 0 | 1 | 0 |
| 29 | 91.84% | 48.81% | 65.56% | 27.36% | 1 | 0 | 1 | 0 |
| 36 | 100.00% | 0.00% | 98.89% | 33.96% | 1 | 0 | 1 | 0 |
| 12 | 100.00% | 21.43% | 98.89% | 11.32% | 1 | 0 | 1 | 0 |
| 7 | 100.00% | 40.48% | 84.44% | 6.60% | 1 | 0 | 1 | 0 |
| | 100.00% | 32.14% | 98.89% | 0.00% | 1 | 0 | 1 | 0 |
| 34 | 100.00% | 0.00% | 98.89% | 32.08% | 1 | 0 | 1 | 0 |
| 33 | 100.00% | 0.00% | 98.89% | 31.13% | 1 | 0 | 1 | 0 |
| | 100.00% | 30.95% | 98.89% | 0.00% | 1 | 0 | 1 | 0 |
| 40 | 92.86% | 0.00% | 98.89% | 37.74% | 1 | 0 | 1 | 0 |
| 18 | 100.00% | 13.10% | 98.89% | 16.98% | 1 | 0 | 1 | 0 |
| 30 | 100.00% | 0.00% | 98.89% | 28.30% | 1 | 0 | 1 | 0 |
| 30 | 100.00% | 0.00% | 98.89% | 28.30% | 1 | 0 | 1 | 0 |
| 29 | 100.00% | 0.00% | 98.89% | 27.36% | 1 | 0 | 1 | 0 |
| 29 | 100.00% | 0.00% | 98.89% | 27.36% | 1 | 0 | 1 | 0 |
| 29 | 100.00% | 0.00% | 98.89% | 27.36% | 1 | 0 | 1 | 0 |
| 25 | 100.00% | 3.57% | 98.89% | 23.58% | 1 | 0 | 1 | 0 |
| 9 | 100.00% | 17.86% | 98.89% | 8.49% | 1 | 0 | 1 | 0 |
| 1 | 100.00% | 25.00% | 98.89% | 0.94% | 1 | 0 | 1 | 0 |
| | 100.00% | 36.90% | 86.67% | 0.00% | 1 | 0 | 1 | 0 |
| 25 | 72.45% | 47.62% | 78.89% | 23.58% | 1 | 0 | 1 | 0 |
| 25 | 100.00% | 0.00% | 98.89% | 23.58% | 1 | 0 | 1 | 0 |
| 25 | 100.00% | 0.00% | 98.89% | 23.58% | 1 | 0 | 1 | 0 |
| 32 | 92.86% | 0.00% | 98.89% | 30.19% | 1 | 0 | 1 | 0 |
| 32 | 92.86% | 0.00% | 98.89% | 30.19% | 1 | 0 | 1 | 0 |
| 32 | 92.86% | 0.00% | 98.89% | 30.19% | 1 | 0 | 1 | 0 |
| 24 | 100.00% | 0.00% | 98.89% | 22.64% | 1 | 0 | 1 | 0 |
| 23 | 95.92% | 69.05% | 34.44% | 21.70% | 1 | 1 | 0 | 0 |
| | 100.00% | 21.43% | 98.89% | 0.00% | 1 | 0 | 1 | 0 |
| 30 | 92.86% | 0.00% | 98.89% | 28.30% | 1 | 0 | 1 | 0 |
| | 100.00% | 20.24% | 98.89% | 0.00% | 1 | 0 | 1 | 0 |
| 23 | 92.86% | 69.05% | 34.44% | 21.70% | 1 | 1 | 0 | 0 |
| | 100.00% | 19.05% | 98.89% | 0.00% | 1 | 0 | 1 | 0 |
| | 100.00% | 19.05% | 98.89% | 0.00% | 1 | 0 | 1 | 0 |
| 29 | 90.82% | 0.00% | 98.89% | 27.36% | 1 | 0 | 1 | 0 |
| | 100.00% | 21.43% | 95.56% | 0.00% | 1 | 0 | 1 | 0 |
| | 100.00% | 17.86% | 98.89% | 0.00% | 1 | 0 | 1 | 0 |
| 5 | 100.00% | 11.90% | 98.89% | 4.72% | 1 | 0 | 1 | 0 |

| | | | | | | | | |
|---|---|---|---|---|---|---|---|---|
| 15 | 100.00% | 2.38% | 98.89% | 14.15% | 1 | 0 | 1 | 0 |
| 17 | 100.00% | 0.00% | 98.89% | 16.04% | 1 | 0 | 1 | 0 |
| 16 | 100.00% | 0.00% | 98.89% | 15.09% | 1 | 0 | 1 | 0 |
| 2 | 100.00% | 28.57% | 83.33% | 1.89% | 1 | 0 | 1 | 0 |
| 15 | 94.90% | 34.52% | 70.00% | 14.15% | 1 | 0 | 1 | 0 |
| 15 | 94.90% | 34.52% | 70.00% | 14.15% | 1 | 0 | 1 | 0 |
| 15 | 100.00% | 0.00% | 98.89% | 14.15% | 1 | 0 | 1 | 0 |
| 15 | 100.00% | 0.00% | 98.89% | 14.15% | 1 | 0 | 1 | 0 |
| 12 | 100.00% | 0.00% | 98.89% | 11.32% | 1 | 0 | 1 | 0 |
| 6 | 87.76% | 82.14% | 33.33% | 5.66% | 1 | 1 | 0 | 0 |
| 10 | 100.00% | 0.00% | 98.89% | 9.43% | 1 | 0 | 1 | 0 |
| 26 | 84.69% | 0.00% | 98.89% | 24.53% | 1 | 0 | 1 | 0 |
| 29 | 81.63% | 0.00% | 98.89% | 27.36% | 1 | 0 | 1 | 0 |
| 9 | 100.00% | 0.00% | 98.89% | 8.49% | 1 | 0 | 1 | 0 |
| 9 | 100.00% | 0.00% | 98.89% | 8.49% | 1 | 0 | 1 | 0 |
| 6 | 100.00% | 97.62% | 3.33% | 5.66% | 1 | 1 | 0 | 0 |
| 8 | 100.00% | 0.00% | 98.89% | 7.55% | 1 | 0 | 1 | 0 |
| | 100.00% | 23.81% | 82.22% | 0.00% | 1 | 0 | 1 | 0 |
| 4 | 100.00% | 0.00% | 98.89% | 3.77% | 1 | 0 | 1 | 0 |
| 3 | 100.00% | 0.00% | 98.89% | 2.83% | 1 | 0 | 1 | 0 |
| | 100.00% | 14.29% | 86.67% | 0.00% | 1 | 0 | 1 | 0 |
| | 100.00% | 1.19% | 98.89% | 0.00% | 1 | 0 | 1 | 0 |
| | 100.00% | 100.00% | 0.00% | 0.00% | 1 | 1 | 0 | 0 |
| | 100.00% | 0.00% | 98.89% | 0.00% | 1 | 0 | 1 | 0 |
| | 100.00% | 0.00% | 98.89% | 0.00% | 1 | 0 | 1 | 0 |
| | 100.00% | 0.00% | 98.89% | 0.00% | 1 | 0 | 1 | 0 |
| | 100.00% | 0.00% | 98.89% | 0.00% | 1 | 0 | 1 | 0 |
| | 100.00% | 0.00% | 98.89% | 0.00% | 1 | 0 | 1 | 0 |
| | 100.00% | 0.00% | 98.89% | 0.00% | 1 | 0 | 1 | 0 |
| | 100.00% | 0.00% | 98.89% | 0.00% | 1 | 0 | 1 | 0 |
| | 100.00% | 0.00% | 98.89% | 0.00% | 1 | 0 | 1 | 0 |
| | 100.00% | 0.00% | 98.89% | 0.00% | 1 | 0 | 1 | 0 |
| | 100.00% | 0.00% | 98.89% | 0.00% | 1 | 0 | 1 | 0 |
| | 100.00% | 0.00% | 98.89% | 0.00% | 1 | 0 | 1 | 0 |
| | 100.00% | 0.00% | 98.89% | 0.00% | 1 | 0 | 1 | 0 |
| | 100.00% | 0.00% | 98.89% | 0.00% | 1 | 0 | 1 | 0 |
| | 100.00% | 0.00% | 98.89% | 0.00% | 1 | 0 | 1 | 0 |
| | 100.00% | 0.00% | 98.89% | 0.00% | 1 | 0 | 1 | 0 |
| | 100.00% | 0.00% | 98.89% | 0.00% | 1 | 0 | 1 | 0 |
| | 100.00% | 0.00% | 98.89% | 0.00% | 1 | 0 | 1 | 0 |
| 31 | 53.06% | 48.81% | 65.56% | 29.25% | 1 | 0 | 1 | 0 |
| | 100.00% | 15.48% | 78.89% | 0.00% | 1 | 0 | 1 | 0 |
| 49 | 50.00% | 47.62% | 47.78% | 46.23% | 0 | 0 | 0 | 0 |

| | | | | | | | | |
|---|---|---|---|---|---|---|---|---|
| 35 | 51.02% | 25.00% | 82.22% | 33.02% | 1 | 0 | 1 | 0 |
| | 97.96% | 0.00% | 91.11% | 0.00% | 1 | 0 | 1 | 0 |
| 2 | 86.73% | 0.00% | 98.89% | 1.89% | 1 | 0 | 1 | 0 |
| | 93.88% | 0.00% | 93.33% | 0.00% | 1 | 0 | 1 | 0 |
| 34 | 74.49% | 19.05% | 61.11% | 32.08% | 1 | 0 | 1 | 0 |
| 34 | 74.49% | 19.05% | 61.11% | 32.08% | 1 | 0 | 1 | 0 |
| 3 | 87.76% | 25.00% | 71.11% | 2.83% | 1 | 0 | 1 | 0 |
| | 100.00% | 44.05% | 42.22% | 0.00% | 1 | 0 | 0 | 0 |
| 26 | 79.59% | 28.57% | 52.22% | 24.53% | 1 | 0 | 1 | 0 |
| | 86.73% | 0.00% | 96.67% | 0.00% | 1 | 0 | 1 | 0 |
| 14 | 77.55% | 5.95% | 85.56% | 13.21% | 1 | 0 | 1 | 0 |
| | 98.98% | 41.67% | 41.11% | 0.00% | 1 | 0 | 0 | 0 |
| | 100.00% | 0.00% | 81.11% | 0.00% | 1 | 0 | 1 | 0 |
| | 81.63% | 0.00% | 98.89% | 0.00% | 1 | 0 | 1 | 0 |
| 9 | 72.45% | 98.81% | 0.00% | 8.49% | 1 | 1 | 0 | 0 |
| 5 | 89.80% | 25.00% | 52.22% | 4.72% | 1 | 0 | 1 | 0 |
| 3 | 74.49% | 0.00% | 92.22% | 2.83% | 1 | 0 | 1 | 0 |
| 17 | 52.04% | 0.00% | 98.89% | 16.04% | 1 | 0 | 1 | 0 |
| 34 | 55.10% | 53.57% | 23.33% | 32.08% | 1 | 1 | 0 | 0 |
| | 86.73% | 0.00% | 76.67% | 0.00% | 1 | 0 | 1 | 0 |
| 31 | 40.82% | 77.38% | 14.44% | 29.25% | 0 | 1 | 0 | 0 |
| | 85.71% | 0.00% | 74.44% | 0.00% | 1 | 0 | 1 | 0 |
| 34 | 60.20% | 33.33% | 34.44% | 32.08% | 1 | 0 | 0 | 0 |
| 1 | 98.98% | 8.33% | 51.11% | 0.94% | 1 | 0 | 1 | 0 |
| 41 | 35.71% | 34.52% | 50.00% | 38.68% | 0 | 0 | 0 | 0 |
| 32 | 54.08% | 39.29% | 30.00% | 30.19% | 1 | 0 | 0 | 0 |
| | 54.08% | 0.00% | 98.89% | 0.00% | 1 | 0 | 1 | 0 |
| 19 | 69.39% | 45.24% | 17.78% | 17.92% | 1 | 0 | 0 | 0 |
| | 51.02% | 0.00% | 98.89% | 0.00% | 1 | 0 | 1 | 0 |
| | 65.31% | 0.00% | 83.33% | 0.00% | 1 | 0 | 1 | 0 |
| | 100.00% | 7.14% | 37.78% | 0.00% | 1 | 0 | 0 | 0 |
| 17 | 39.80% | 0.00% | 88.89% | 16.04% | 0 | 0 | 1 | 0 |
| | 100.00% | 0.00% | 41.11% | 0.00% | 1 | 0 | 0 | 0 |
| 9 | 44.90% | 0.00% | 85.56% | 8.49% | 0 | 0 | 1 | 0 |
| 25 | 50.00% | 16.67% | 45.56% | 23.58% | 0 | 0 | 0 | 0 |
| | 100.00% | 0.00% | 34.44% | 0.00% | 1 | 0 | 0 | 0 |
| 16 | 69.39% | 25.00% | 23.33% | 15.09% | 1 | 0 | 0 | 0 |
| | 32.65% | 0.00% | 98.89% | 0.00% | 0 | 0 | 1 | 0 |
| 11 | 65.31% | 40.48% | 14.44% | 10.38% | 1 | 0 | 0 | 0 |
| 9 | 63.27% | 35.71% | 22.22% | 8.49% | 1 | 0 | 0 | 0 |
| 9 | 66.33% | 0.00% | 53.33% | 8.49% | 1 | 0 | 1 | 0 |
| | 22.45% | 0.00% | 98.89% | 0.00% | 0 | 0 | 1 | 0 |
| 15 | 28.57% | 0.00% | 77.78% | 14.15% | 0 | 0 | 1 | 0 |

| | | | | | | | | |
|---|---|---|---|---|---|---|---|---|
| | 65.31% | 53.57% | 0.00% | 0.00% | 1 | 1 | 0 | 0 |
| 18 | 47.96% | 42.86% | 10.00% | 16.98% | 0 | 0 | 0 | 0 |
| | 18.37% | 0.00% | 97.78% | 0.00% | 0 | 0 | 1 | 0 |
| 9 | 65.31% | 30.95% | 11.11% | 8.49% | 1 | 0 | 0 | 0 |
| 9 | 57.14% | 0.00% | 50.00% | 8.49% | 1 | 0 | 0 | 0 |
| 23 | 16.33% | 0.00% | 75.56% | 21.70% | 0 | 0 | 1 | 0 |
| 23 | 19.39% | 0.00% | 71.11% | 21.70% | 0 | 0 | 1 | 0 |
| 10 | 39.80% | 0.00% | 62.22% | 9.43% | 0 | 0 | 1 | 0 |
| 30 | 8.16% | 0.00% | 74.44% | 28.30% | 0 | 0 | 1 | 0 |
| | 13.27% | 0.00% | 96.67% | 0.00% | 0 | 0 | 1 | 0 |
| | 39.80% | 48.81% | 20.00% | 0.00% | 0 | 0 | 0 | 0 |
| 1 | 53.06% | 0.00% | 51.11% | 0.94% | 1 | 0 | 1 | 0 |
| 4 | 37.76% | 63.10% | 0.00% | 3.77% | 0 | 1 | 0 | 0 |
| 4 | 37.76% | 63.10% | 0.00% | 3.77% | 0 | 1 | 0 | 0 |
| | 39.80% | 48.81% | 14.44% | 0.00% | 0 | 0 | 0 | 0 |
| | 42.86% | 0.00% | 60.00% | 0.00% | 0 | 0 | 1 | 0 |
| | 81.63% | 0.00% | 20.00% | 0.00% | 1 | 0 | 0 | 0 |
| | 100.00% | 0.00% | 0.00% | 0.00% | 1 | 0 | 0 | 0 |
| 9 | 51.02% | 23.81% | 15.56% | 8.49% | 1 | 0 | 0 | 0 |
| | 0.00% | 98.81% | 0.00% | 0.00% | 0 | 1 | 0 | 0 |
| 28 | 2.04% | 0.00% | 70.00% | 26.42% | 0 | 0 | 1 | 0 |
| | 41.84% | 0.00% | 55.56% | 0.00% | 0 | 0 | 1 | 0 |
| 4 | 5.10% | 0.00% | 87.78% | 3.77% | 0 | 0 | 1 | 0 |
| 49 | 0.00% | 2.38% | 47.78% | 46.23% | 0 | 0 | 0 | 0 |
| 49 | 0.00% | 2.38% | 47.78% | 46.23% | 0 | 0 | 0 | 0 |
| 7 | 43.88% | 45.24% | 0.00% | 6.60% | 0 | 0 | 0 | 0 |
| 6 | 43.88% | 45.24% | 0.00% | 5.66% | 0 | 0 | 0 | 0 |
| 9 | 42.86% | 10.71% | 32.22% | 8.49% | 0 | 0 | 0 | 0 |
| | 60.20% | 2.38% | 31.11% | 0.00% | 1 | 0 | 0 | 0 |
| | 60.20% | 2.38% | 31.11% | 0.00% | 1 | 0 | 0 | 0 |
| 6 | 43.88% | 44.05% | 0.00% | 5.66% | 0 | 0 | 0 | 0 |
| | 53.06% | 0.00% | 38.89% | 0.00% | 1 | 0 | 0 | 0 |
| 13 | 43.88% | 35.71% | 0.00% | 12.26% | 0 | 0 | 0 | 0 |
| 3 | 32.65% | 0.00% | 55.56% | 2.83% | 0 | 0 | 1 | 0 |
| 22 | 0.00% | 0.00% | 70.00% | 20.75% | 0 | 0 | 1 | 0 |
| | 10.20% | 0.00% | 80.00% | 0.00% | 0 | 0 | 1 | 0 |
| 6 | 43.88% | 40.48% | 0.00% | 5.66% | 0 | 0 | 0 | 0 |
| | 89.80% | 0.00% | 0.00% | 0.00% | 1 | 0 | 0 | 0 |
| | 32.65% | 57.14% | 0.00% | 0.00% | 0 | 1 | 0 | 0 |
| 49 | 0.00% | 0.00% | 43.33% | 46.23% | 0 | 0 | 0 | 0 |
| | 39.80% | 48.81% | 0.00% | 0.00% | 0 | 0 | 0 | 0 |
| 49 | 0.00% | 0.00% | 42.22% | 46.23% | 0 | 0 | 0 | 0 |
| 43 | 0.00% | 0.00% | 47.78% | 40.57% | 0 | 0 | 0 | 0 |

| | | | | | | | | |
|---|---|---|---|---|---|---|---|---|
| | 53.06% | 0.00% | 34.44% | 0.00% | 1 | 0 | 0 | 0 |
| | 0.00% | 0.00% | 86.67% | 0.00% | 0 | 0 | 1 | 0 |
| 2 | 46.94% | 0.00% | 37.78% | 1.89% | 0 | 0 | 0 | 0 |
| 35 | 11.22% | 32.14% | 10.00% | 33.02% | 0 | 0 | 0 | 0 |
| 6 | 43.88% | 35.71% | 0.00% | 5.66% | 0 | 0 | 0 | 0 |
| 2 | 31.63% | 51.19% | 0.00% | 1.89% | 0 | 1 | 0 | 0 |
| 12 | 37.76% | 14.29% | 21.11% | 11.32% | 0 | 0 | 0 | 0 |
| 15 | 35.71% | 34.52% | 0.00% | 14.15% | 0 | 0 | 0 | 0 |
| | 36.73% | 47.62% | 0.00% | 0.00% | 0 | 0 | 0 | 0 |
| 5 | 43.88% | 35.71% | 0.00% | 4.72% | 0 | 0 | 0 | 0 |
| | 8.16% | 0.00% | 75.56% | 0.00% | 0 | 0 | 1 | 0 |
| | 34.69% | 0.00% | 48.89% | 0.00% | 0 | 0 | 0 | 0 |
| | 29.59% | 0.00% | 53.33% | 0.00% | 0 | 0 | 1 | 0 |
| 5 | 41.84% | 35.71% | 0.00% | 4.72% | 0 | 0 | 0 | 0 |
| 5 | 41.84% | 35.71% | 0.00% | 4.72% | 0 | 0 | 0 | 0 |
| | 47.96% | 14.29% | 20.00% | 0.00% | 0 | 0 | 0 | 0 |
| 7 | 46.94% | 11.90% | 16.67% | 6.60% | 0 | 0 | 0 | 0 |
| 12 | 9.18% | 0.00% | 61.11% | 11.32% | 0 | 0 | 1 | 0 |
| 2 | 43.88% | 35.71% | 0.00% | 1.89% | 0 | 0 | 0 | 0 |
| 2 | 43.88% | 35.71% | 0.00% | 1.89% | 0 | 0 | 0 | 0 |
| 39 | 7.14% | 35.71% | 0.00% | 36.79% | 0 | 0 | 0 | 0 |
| 14 | 7.14% | 0.00% | 56.67% | 13.21% | 0 | 0 | 1 | 0 |
| 4 | 27.55% | 0.00% | 45.56% | 3.77% | 0 | 0 | 0 | 0 |
| 4 | 27.55% | 0.00% | 45.56% | 3.77% | 0 | 0 | 0 | 0 |
| | 50.00% | 1.19% | 25.56% | 0.00% | 0 | 0 | 0 | 0 |
| 32 | 13.27% | 32.14% | 0.00% | 30.19% | 0 | 0 | 0 | 0 |
| | 39.80% | 35.71% | 0.00% | 0.00% | 0 | 0 | 0 | 0 |
| | 61.22% | 14.29% | 0.00% | 0.00% | 1 | 0 | 0 | 0 |
| 5 | 35.71% | 34.52% | 0.00% | 4.72% | 0 | 0 | 0 | 0 |
| | 40.82% | 33.33% | 0.00% | 0.00% | 0 | 0 | 0 | 0 |
| | 9.18% | 0.00% | 64.44% | 0.00% | 0 | 0 | 1 | 0 |
| | 9.18% | 64.29% | 0.00% | 0.00% | 0 | 1 | 0 | 0 |
| 12 | 7.14% | 0.00% | 54.44% | 11.32% | 0 | 0 | 1 | 0 |
| 2 | 1.02% | 0.00% | 70.00% | 1.89% | 0 | 0 | 1 | 0 |
| | 35.71% | 36.90% | 0.00% | 0.00% | 0 | 0 | 0 | 0 |
| | 0.00% | 71.43% | 0.00% | 0.00% | 0 | 1 | 0 | 0 |
| 3 | 0.00% | 0.00% | 67.78% | 2.83% | 0 | 0 | 1 | 0 |
| 9 | 31.63% | 15.48% | 14.44% | 8.49% | 0 | 0 | 0 | 0 |
| 4 | 17.35% | 48.81% | 0.00% | 3.77% | 0 | 0 | 0 | 0 |
| | 5.10% | 0.00% | 64.44% | 0.00% | 0 | 0 | 1 | 0 |
| 2 | 30.61% | 34.52% | 0.00% | 1.89% | 0 | 0 | 0 | 0 |
| 2 | 0.00% | 0.00% | 64.44% | 1.89% | 0 | 0 | 1 | 0 |
| 6 | 15.31% | 0.00% | 44.44% | 5.66% | 0 | 0 | 0 | 0 |

| | | | | | | | | |
|---|---|---|---|---|---|---|---|---|
| 2 | 0.00% | 0.00% | 63.33% | 1.89% | 0 | 0 | 1 | 0 |
| 9 | 22.45% | 33.33% | 0.00% | 8.49% | 0 | 0 | 0 | 0 |
| | 39.80% | 0.00% | 24.44% | 0.00% | 0 | 0 | 0 | 0 |
| 18 | 10.20% | 20.24% | 16.67% | 16.98% | 0 | 0 | 0 | 0 |
| | 0.00% | 0.00% | 63.33% | 0.00% | 0 | 0 | 1 | 0 |
| 2 | 37.76% | 14.29% | 8.89% | 1.89% | 0 | 0 | 0 | 0 |
| | 0.00% | 61.90% | 0.00% | 0.00% | 0 | 1 | 0 | 0 |
| 49 | 0.00% | 0.00% | 15.56% | 46.23% | 0 | 0 | 0 | 0 |
| 24 | 5.10% | 0.00% | 32.22% | 22.64% | 0 | 0 | 0 | 0 |
| | 27.55% | 32.14% | 0.00% | 0.00% | 0 | 0 | 0 | 0 |
| 50 | 0.00% | 2.38% | 10.00% | 47.17% | 0 | 0 | 0 | 0 |
| | 0.00% | 0.00% | 58.89% | 0.00% | 0 | 0 | 1 | 0 |
| | 0.00% | 0.00% | 58.89% | 0.00% | 0 | 0 | 1 | 0 |
| | 53.06% | 0.00% | 5.56% | 0.00% | 1 | 0 | 0 | 0 |
| | 53.06% | 0.00% | 4.44% | 0.00% | 1 | 0 | 0 | 0 |
| | 47.96% | 0.00% | 8.89% | 0.00% | 0 | 0 | 0 | 0 |
| | 2.04% | 0.00% | 54.44% | 0.00% | 0 | 0 | 1 | 0 |
| | 29.59% | 0.00% | 26.67% | 0.00% | 0 | 0 | 0 | 0 |
| | 56.12% | 0.00% | 0.00% | 0.00% | 1 | 0 | 0 | 0 |
| | 53.06% | 0.00% | 2.22% | 0.00% | 1 | 0 | 0 | 0 |
| 8 | 26.53% | 9.52% | 10.00% | 7.55% | 0 | 0 | 0 | 0 |
| | 47.96% | 0.00% | 5.56% | 0.00% | 0 | 0 | 0 | 0 |
| | 53.06% | 0.00% | 0.00% | 0.00% | 1 | 0 | 0 | 0 |
| 18 | 0.00% | 35.71% | 0.00% | 16.98% | 0 | 0 | 0 | 0 |
| 8 | 26.53% | 11.90% | 6.67% | 7.55% | 0 | 0 | 0 | 0 |
| | 52.04% | 0.00% | 0.00% | 0.00% | 1 | 0 | 0 | 0 |
| 53 | 0.00% | 0.00% | 1.11% | 50.00% | 0 | 0 | 0 | 0 |
| 53 | 0.00% | 0.00% | 1.11% | 50.00% | 0 | 0 | 0 | 0 |
| 53 | 0.00% | 0.00% | 1.11% | 50.00% | 0 | 0 | 0 | 0 |
| 53 | 0.00% | 0.00% | 1.11% | 50.00% | 0 | 0 | 0 | 0 |
| 24 | 0.00% | 0.00% | 26.67% | 22.64% | 0 | 0 | 0 | 0 |
| 4 | 1.02% | 0.00% | 44.44% | 3.77% | 0 | 0 | 0 | 0 |
| 2 | 0.00% | 0.00% | 45.56% | 1.89% | 0 | 0 | 0 | 0 |
| 45 | 0.00% | 0.00% | 4.44% | 42.45% | 0 | 0 | 0 | 0 |
| | 37.76% | 0.00% | 8.89% | 0.00% | 0 | 0 | 0 | 0 |
| | 21.43% | 25.00% | 0.00% | 0.00% | 0 | 0 | 0 | 0 |
| 2 | 0.00% | 0.00% | 43.33% | 1.89% | 0 | 0 | 0 | 0 |
| 15 | 13.27% | 15.48% | 2.22% | 14.15% | 0 | 0 | 0 | 0 |
| 18 | 0.00% | 0.00% | 27.78% | 16.98% | 0 | 0 | 0 | 0 |
| 2 | 0.00% | 0.00% | 42.22% | 1.89% | 0 | 0 | 0 | 0 |
| 2 | 0.00% | 0.00% | 42.22% | 1.89% | 0 | 0 | 0 | 0 |
| 22 | 0.00% | 0.00% | 23.33% | 20.75% | 0 | 0 | 0 | 0 |
| | 0.00% | 0.00% | 43.33% | 0.00% | 0 | 0 | 0 | 0 |

| | | | | | | | | |
|---|---|---|---|---|---|---|---|---|
| | 0.00% | 0.00% | 43.33% | 0.00% | 0 | 0 | 0 | 0 |
| | 0.00% | 0.00% | 43.33% | 0.00% | 0 | 0 | 0 | 0 |
| 20 | 0.00% | 0.00% | 24.44% | 18.87% | 0 | 0 | 0 | 0 |
| | 40.82% | 0.00% | 0.00% | 0.00% | 0 | 0 | 0 | 0 |
| 26 | 0.00% | 0.00% | 15.56% | 24.53% | 0 | 0 | 0 | 0 |
| 2 | 0.00% | 0.00% | 37.78% | 1.89% | 0 | 0 | 0 | 0 |
| 5 | 21.43% | 11.90% | 1.11% | 4.72% | 0 | 0 | 0 | 0 |
| 9 | 13.27% | 9.52% | 6.67% | 8.49% | 0 | 0 | 0 | 0 |
| 20 | 0.00% | 0.00% | 18.89% | 18.87% | 0 | 0 | 0 | 0 |
| | 26.53% | 1.19% | 10.00% | 0.00% | 0 | 0 | 0 | 0 |
| 36 | 0.00% | 3.57% | 0.00% | 33.96% | 0 | 0 | 0 | 0 |
| | 17.35% | 0.00% | 20.00% | 0.00% | 0 | 0 | 0 | 0 |
| 1 | 16.33% | 5.95% | 13.33% | 0.94% | 0 | 0 | 0 | 0 |
| | 14.29% | 0.00% | 22.22% | 0.00% | 0 | 0 | 0 | 0 |
| 2 | 0.00% | 0.00% | 34.44% | 1.89% | 0 | 0 | 0 | 0 |
| 9 | 5.10% | 0.00% | 22.22% | 8.49% | 0 | 0 | 0 | 0 |
| | 34.69% | 0.00% | 1.11% | 0.00% | 0 | 0 | 0 | 0 |
| | 27.55% | 0.00% | 6.67% | 0.00% | 0 | 0 | 0 | 0 |
| | 28.57% | 0.00% | 5.56% | 0.00% | 0 | 0 | 0 | 0 |
| 2 | 0.00% | 0.00% | 32.22% | 1.89% | 0 | 0 | 0 | 0 |
| 22 | 0.00% | 0.00% | 12.22% | 20.75% | 0 | 0 | 0 | 0 |
| | 30.61% | 0.00% | 2.22% | 0.00% | 0 | 0 | 0 | 0 |
| 8 | 10.20% | 8.33% | 6.67% | 7.55% | 0 | 0 | 0 | 0 |
| 3 | 15.31% | 14.29% | 0.00% | 2.83% | 0 | 0 | 0 | 0 |
| 2 | 0.00% | 0.00% | 30.00% | 1.89% | 0 | 0 | 0 | 0 |
| 9 | 0.00% | 0.00% | 23.33% | 8.49% | 0 | 0 | 0 | 0 |
| | 5.10% | 26.19% | 0.00% | 0.00% | 0 | 0 | 0 | 0 |
| | 29.59% | 0.00% | 0.00% | 0.00% | 0 | 0 | 0 | 0 |
| | 17.35% | 0.00% | 12.22% | 0.00% | 0 | 0 | 0 | 0 |
| | 7.14% | 0.00% | 22.22% | 0.00% | 0 | 0 | 0 | 0 |
| | 0.00% | 0.00% | 28.89% | 0.00% | 0 | 0 | 0 | 0 |
| | 22.45% | 0.00% | 4.44% | 0.00% | 0 | 0 | 0 | 0 |
| | 12.24% | 0.00% | 13.33% | 0.00% | 0 | 0 | 0 | 0 |
| | 0.00% | 0.00% | 25.56% | 0.00% | 0 | 0 | 0 | 0 |
| | 0.00% | 0.00% | 24.44% | 0.00% | 0 | 0 | 0 | 0 |
| 2 | 0.00% | 0.00% | 22.22% | 1.89% | 0 | 0 | 0 | 0 |
| | 0.00% | 0.00% | 23.33% | 0.00% | 0 | 0 | 0 | 0 |
| 19 | 0.00% | 4.76% | 0.00% | 17.92% | 0 | 0 | 0 | 0 |
| | 22.45% | 0.00% | 0.00% | 0.00% | 0 | 0 | 0 | 0 |
| | 22.45% | 0.00% | 0.00% | 0.00% | 0 | 0 | 0 | 0 |
| | 0.00% | 0.00% | 22.22% | 0.00% | 0 | 0 | 0 | 0 |
| 7 | 0.00% | 0.00% | 15.56% | 6.60% | 0 | 0 | 0 | 0 |
| 23 | 0.00% | 0.00% | 0.00% | 21.70% | 0 | 0 | 0 | 0 |

| | | | | | | | | |
|---|---|---|---|---|---|---|---|---|
| 23 | 0.00% | 0.00% | 0.00% | 21.70% | 0 | 0 | 0 | 0 |
| 23 | 0.00% | 0.00% | 0.00% | 21.70% | 0 | 0 | 0 | 0 |
| | 21.43% | 0.00% | 0.00% | 0.00% | 0 | 0 | 0 | 0 |
| 7 | 0.00% | 0.00% | 14.44% | 6.60% | 0 | 0 | 0 | 0 |
| 14 | 0.00% | 0.00% | 7.78% | 13.21% | 0 | 0 | 0 | 0 |
| 22 | 0.00% | 0.00% | 0.00% | 20.75% | 0 | 0 | 0 | 0 |
| | 0.00% | 0.00% | 20.00% | 0.00% | 0 | 0 | 0 | 0 |
| | 0.00% | 19.05% | 0.00% | 0.00% | 0 | 0 | 0 | 0 |
| | 0.00% | 19.05% | 0.00% | 0.00% | 0 | 0 | 0 | 0 |
| | 11.22% | 0.00% | 7.78% | 0.00% | 0 | 0 | 0 | 0 |
| | 0.00% | 0.00% | 18.89% | 0.00% | 0 | 0 | 0 | 0 |
| | 9.18% | 0.00% | 8.89% | 0.00% | 0 | 0 | 0 | 0 |
| | 0.00% | 0.00% | 17.78% | 0.00% | 0 | 0 | 0 | 0 |
| | 0.00% | 0.00% | 17.78% | 0.00% | 0 | 0 | 0 | 0 |
| | 0.00% | 0.00% | 17.78% | 0.00% | 0 | 0 | 0 | 0 |
| | 2.04% | 0.00% | 15.56% | 0.00% | 0 | 0 | 0 | 0 |
| | 15.31% | 0.00% | 2.22% | 0.00% | 0 | 0 | 0 | 0 |
| 12 | 0.00% | 4.76% | 0.00% | 11.32% | 0 | 0 | 0 | 0 |
| | 0.00% | 0.00% | 15.56% | 0.00% | 0 | 0 | 0 | 0 |
| | 0.00% | 0.00% | 15.56% | 0.00% | 0 | 0 | 0 | 0 |
| | 0.00% | 0.00% | 14.44% | 0.00% | 0 | 0 | 0 | 0 |
| | 0.00% | 0.00% | 14.44% | 0.00% | 0 | 0 | 0 | 0 |
| | 14.29% | 0.00% | 0.00% | 0.00% | 0 | 0 | 0 | 0 |
| 15 | 0.00% | 0.00% | 0.00% | 14.15% | 0 | 0 | 0 | 0 |
| 15 | 0.00% | 0.00% | 0.00% | 14.15% | 0 | 0 | 0 | 0 |
| 15 | 0.00% | 0.00% | 0.00% | 14.15% | 0 | 0 | 0 | 0 |
| | 0.00% | 0.00% | 13.33% | 0.00% | 0 | 0 | 0 | 0 |
| | 0.00% | 0.00% | 13.33% | 0.00% | 0 | 0 | 0 | 0 |
| 13 | 0.00% | 0.00% | 0.00% | 12.26% | 0 | 0 | 0 | 0 |
| | 0.00% | 11.90% | 0.00% | 0.00% | 0 | 0 | 0 | 0 |
| | 0.00% | 11.90% | 0.00% | 0.00% | 0 | 0 | 0 | 0 |
| 12 | 0.00% | 0.00% | 0.00% | 11.32% | 0 | 0 | 0 | 0 |
| 12 | 0.00% | 0.00% | 0.00% | 11.32% | 0 | 0 | 0 | 0 |
| 12 | 0.00% | 0.00% | 0.00% | 11.32% | 0 | 0 | 0 | 0 |
| 12 | 0.00% | 0.00% | 0.00% | 11.32% | 0 | 0 | 0 | 0 |
| 12 | 0.00% | 0.00% | 0.00% | 11.32% | 0 | 0 | 0 | 0 |
| | 11.22% | 0.00% | 0.00% | 0.00% | 0 | 0 | 0 | 0 |
| | 0.00% | 0.00% | 11.11% | 0.00% | 0 | 0 | 0 | 0 |
| | 0.00% | 10.71% | 0.00% | 0.00% | 0 | 0 | 0 | 0 |
| | 0.00% | 0.00% | 10.00% | 0.00% | 0 | 0 | 0 | 0 |
| | 0.00% | 9.52% | 0.00% | 0.00% | 0 | 0 | 0 | 0 |
| 10 | 0.00% | 0.00% | 0.00% | 9.43% | 0 | 0 | 0 | 0 |
| | 9.18% | 0.00% | 0.00% | 0.00% | 0 | 0 | 0 | 0 |

| | | | | | | | | |
|---|---|---|---|---|---|---|---|---|
| 2 | 0.00% | 7.14% | 0.00% | 1.89% | 0 | 0 | 0 | 0 |
| | 0.00% | 0.00% | 8.89% | 0.00% | 0 | 0 | 0 | 0 |
| | 0.00% | 0.00% | 8.89% | 0.00% | 0 | 0 | 0 | 0 |
| | 0.00% | 0.00% | 8.89% | 0.00% | 0 | 0 | 0 | 0 |
| | 0.00% | 0.00% | 8.89% | 0.00% | 0 | 0 | 0 | 0 |
| 9 | 0.00% | 0.00% | 0.00% | 8.49% | 0 | 0 | 0 | 0 |
| 9 | 0.00% | 0.00% | 0.00% | 8.49% | 0 | 0 | 0 | 0 |
| 9 | 0.00% | 0.00% | 0.00% | 8.49% | 0 | 0 | 0 | 0 |
| | 0.00% | 8.33% | 0.00% | 0.00% | 0 | 0 | 0 | 0 |
| | 0.00% | 8.33% | 0.00% | 0.00% | 0 | 0 | 0 | 0 |
| 7 | 0.00% | 1.19% | 0.00% | 6.60% | 0 | 0 | 0 | 0 |
| 7 | 0.00% | 1.19% | 0.00% | 6.60% | 0 | 0 | 0 | 0 |
| | 0.00% | 0.00% | 7.78% | 0.00% | 0 | 0 | 0 | 0 |
| 8 | 0.00% | 0.00% | 0.00% | 7.55% | 0 | 0 | 0 | 0 |
| 8 | 0.00% | 0.00% | 0.00% | 7.55% | 0 | 0 | 0 | 0 |
| 8 | 0.00% | 0.00% | 0.00% | 7.55% | 0 | 0 | 0 | 0 |
| 8 | 0.00% | 0.00% | 0.00% | 7.55% | 0 | 0 | 0 | 0 |
| | 0.00% | 7.14% | 0.00% | 0.00% | 0 | 0 | 0 | 0 |
| 7 | 0.00% | 0.00% | 0.00% | 6.60% | 0 | 0 | 0 | 0 |
| 7 | 0.00% | 0.00% | 0.00% | 6.60% | 0 | 0 | 0 | 0 |
| 7 | 0.00% | 0.00% | 0.00% | 6.60% | 0 | 0 | 0 | 0 |
| 7 | 0.00% | 0.00% | 0.00% | 6.60% | 0 | 0 | 0 | 0 |
| 7 | 0.00% | 0.00% | 0.00% | 6.60% | 0 | 0 | 0 | 0 |
| | 6.12% | 0.00% | 0.00% | 0.00% | 0 | 0 | 0 | 0 |
| | 0.00% | 5.95% | 0.00% | 0.00% | 0 | 0 | 0 | 0 |
| | 0.00% | 5.95% | 0.00% | 0.00% | 0 | 0 | 0 | 0 |
| | 0.00% | 5.95% | 0.00% | 0.00% | 0 | 0 | 0 | 0 |
| 6 | 0.00% | 0.00% | 0.00% | 5.66% | 0 | 0 | 0 | 0 |
| 6 | 0.00% | 0.00% | 0.00% | 5.66% | 0 | 0 | 0 | 0 |
| 6 | 0.00% | 0.00% | 0.00% | 5.66% | 0 | 0 | 0 | 0 |
| 6 | 0.00% | 0.00% | 0.00% | 5.66% | 0 | 0 | 0 | 0 |
| 6 | 0.00% | 0.00% | 0.00% | 5.66% | 0 | 0 | 0 | 0 |
| 6 | 0.00% | 0.00% | 0.00% | 5.66% | 0 | 0 | 0 | 0 |
| 6 | 0.00% | 0.00% | 0.00% | 5.66% | 0 | 0 | 0 | 0 |
| 6 | 0.00% | 0.00% | 0.00% | 5.66% | 0 | 0 | 0 | 0 |
| | 0.00% | 0.00% | 5.56% | 0.00% | 0 | 0 | 0 | 0 |
| | 0.00% | 0.00% | 5.56% | 0.00% | 0 | 0 | 0 | 0 |
| | 5.10% | 0.00% | 0.00% | 0.00% | 0 | 0 | 0 | 0 |
| 1 | 4.08% | 0.00% | 0.00% | 0.94% | 0 | 0 | 0 | 0 |
| | 0.00% | 4.76% | 0.00% | 0.00% | 0 | 0 | 0 | 0 |
| 5 | 0.00% | 0.00% | 0.00% | 4.72% | 0 | 0 | 0 | 0 |
| | 0.00% | 0.00% | 4.44% | 0.00% | 0 | 0 | 0 | 0 |
| | 0.00% | 0.00% | 4.44% | 0.00% | 0 | 0 | 0 | 0 |

| | | | | | | | | |
|---|---|---|---|---|---|---|---|---|
| 4 | 0.00% | 0.00% | 0.00% | 3.77% | 0 | 0 | 0 | 0 |
| 4 | 0.00% | 0.00% | 0.00% | 3.77% | 0 | 0 | 0 | 0 |
| 4 | 0.00% | 0.00% | 0.00% | 3.77% | 0 | 0 | 0 | 0 |
| 4 | 0.00% | 0.00% | 0.00% | 3.77% | 0 | 0 | 0 | 0 |
| 4 | 0.00% | 0.00% | 0.00% | 3.77% | 0 | 0 | 0 | 0 |
| 4 | 0.00% | 0.00% | 0.00% | 3.77% | 0 | 0 | 0 | 0 |
| | 0.00% | 3.57% | 0.00% | 0.00% | 0 | 0 | 0 | 0 |
| | 0.00% | 0.00% | 3.33% | 0.00% | 0 | 0 | 0 | 0 |
| | 0.00% | 0.00% | 3.33% | 0.00% | 0 | 0 | 0 | 0 |
| | 0.00% | 0.00% | 3.33% | 0.00% | 0 | 0 | 0 | 0 |
| | 0.00% | 0.00% | 3.33% | 0.00% | 0 | 0 | 0 | 0 |
| | 0.00% | 0.00% | 3.33% | 0.00% | 0 | 0 | 0 | 0 |
| | 3.06% | 0.00% | 0.00% | 0.00% | 0 | 0 | 0 | 0 |
| 3 | 0.00% | 0.00% | 0.00% | 2.83% | 0 | 0 | 0 | 0 |
| 3 | 0.00% | 0.00% | 0.00% | 2.83% | 0 | 0 | 0 | 0 |
| 3 | 0.00% | 0.00% | 0.00% | 2.83% | 0 | 0 | 0 | 0 |
| 3 | 0.00% | 0.00% | 0.00% | 2.83% | 0 | 0 | 0 | 0 |
| 3 | 0.00% | 0.00% | 0.00% | 2.83% | 0 | 0 | 0 | 0 |
| | 0.00% | 2.38% | 0.00% | 0.00% | 0 | 0 | 0 | 0 |
| | 0.00% | 0.00% | 2.22% | 0.00% | 0 | 0 | 0 | 0 |
| | 0.00% | 0.00% | 2.22% | 0.00% | 0 | 0 | 0 | 0 |
| | 2.04% | 0.00% | 0.00% | 0.00% | 0 | 0 | 0 | 0 |
| | 2.04% | 0.00% | 0.00% | 0.00% | 0 | 0 | 0 | 0 |
| | 2.04% | 0.00% | 0.00% | 0.00% | 0 | 0 | 0 | 0 |
| 2 | 0.00% | 0.00% | 0.00% | 1.89% | 0 | 0 | 0 | 0 |
| 2 | 0.00% | 0.00% | 0.00% | 1.89% | 0 | 0 | 0 | 0 |
| 2 | 0.00% | 0.00% | 0.00% | 1.89% | 0 | 0 | 0 | 0 |
| 2 | 0.00% | 0.00% | 0.00% | 1.89% | 0 | 0 | 0 | 0 |
| | 0.00% | 1.19% | 0.00% | 0.00% | 0 | 0 | 0 | 0 |
| | 0.00% | 1.19% | 0.00% | 0.00% | 0 | 0 | 0 | 0 |
| | 0.00% | 1.19% | 0.00% | 0.00% | 0 | 0 | 0 | 0 |
| | 0.00% | 0.00% | 1.11% | 0.00% | 0 | 0 | 0 | 0 |
| | 0.00% | 0.00% | 1.11% | 0.00% | 0 | 0 | 0 | 0 |
| | 1.02% | 0.00% | 0.00% | 0.00% | 0 | 0 | 0 | 0 |
| 1 | 0.00% | 0.00% | 0.00% | 0.94% | 0 | 0 | 0 | 0 |
| 1 | 0.00% | 0.00% | 0.00% | 0.94% | 0 | 0 | 0 | 0 |
| 1 | 0.00% | 0.00% | 0.00% | 0.94% | 0 | 0 | 0 | 0 |
| 1 | 0.00% | 0.00% | 0.00% | 0.94% | 0 | 0 | 0 | 0 |
| 1 | 0.00% | 0.00% | 0.00% | 0.94% | 0 | 0 | 0 | 0 |
| 1 | 0.00% | 0.00% | 0.00% | 0.94% | 0 | 0 | 0 | 0 |
| 1 | 0.00% | 0.00% | 0.00% | 0.94% | 0 | 0 | 0 | 0 |
