## Supplemental table 4 for "A systems genomics approach to uncover patient-specific pathogenic pathways and proteins in a complex disease"

Inflamed patients

| Cluster | Platform | All Patinets | LSP1 + | HDAC7 + | MAML2 + | DNMT3B + | Female |
|---|---|---|---|---|---|---|---|
| **1** | Human Genome U133 Plus 2.0 | 1 | 0 | 0 | 0 | 1 | 0 |
| | Affymetrix GeneChip Human Gene 1.0 | 7 | 3 | 5 | 2 | 2 | 3 |
| **2** | Human Genome U133 Plus 2.0 | 5 | 2 | 4 | 3 | 3 | 3 |
| | Affymetrix GeneChip Human Gene 1.0 | 8 | 4 | 5 | 4 | 5 | 4 |
| **3** | Human Genome U133 Plus 2.0 | 12 | 5 | 8 | 8 | 5 | 4 |
| | Affymetrix GeneChip Human Gene 1.0 | 4 | 1 | 1 | 1 | 4 | 2 |
| **4** | Human Genome U133 Plus 2.0 | 4 | 1 | 2 | 1 | 1 | 2 |
| | Affymetrix GeneChip Human Gene 1.0 | 16 | 5 | 11 | 4 | 7 | 5 |
| **All** | Human Genome U133 Plus 2.0 | 22 | 8 | 14 | 12 | 10 | 9 |
| | Affymetrix GeneChip Human Gene 1.0 | 35 | 13 | 22 | 11 | 18 | 14 |

Non-inflamed patients

| Cluster | Platform | All Patinets | LSP1 + | HDAC7 + | MAML2 + | DNMT3B + | Female |
|---|---|---|---|---|---|---|---|
| **1** | Affymetrix GeneChip Human Gene 1.0 | 5 | 2 | 3 | 2 | 2 | 1 |
| **2** | Affymetrix GeneChip Human Gene 1.0 | 12 | 7 | 6 | 7 | 7 | 9 |
| **3** | Affymetrix GeneChip Human Gene 1.0 | 7 | 2 | 2 | 4 | 3 | 4 |
| **4** | Affymetrix GeneChip Human Gene 1.0 | 16 | 8 | 14 | 5 | 7 | 8 |
| **All** | | 40 | 19 | 25 | 18 | 19 | 22 |

| Male | Age mean (range) | Age at diagnosis mena (range) |
|---|---|---|
| 1 | 48 (48) | 29 (29) |
| 4 | 42.1 (29 -63) | 31.8 (11-58) |
| 2 | 44.6 (27-76) | 25.2 (19-35) |
| 4 | 44.1 (22-59) | 31.8 (16-58) |
| 8 | 52.8 (31-73) | 37.8 (9-64) |
| 2 | 48 (34-59) | 37 (24-52) |
| 2 | 48.7 (37-56) | 30.8 (12-47) |
| 7 | 40.1 (24-52) | 27 (14-40) |
| 13 | | |
| 17 | | |

| Male | Age mean (range) | Age at diagnosis mena (range) |
|---|---|---|
| 4 | 50.2 (29-67) | 40.2 (22-64) |
| 3 | 39 (22-65) | 27.3 (16-58) |
| 3 | 44 (31-66) | 23.1 (9-51) |
| 8 | 48.6 (32-77) | 35 (12-66) |
| 18 | | |
