## Supplemental table 5 for "A systems genomics approach to uncover patient-specific pathogenic pathways and proteins in a complex disease"

| SNP_ID | ID of SNP affected gene | Name of affected gene | Type of regulatory effect | iant componenet of the | nvovled in UK IBD datase |
|---|---|---|---|---|---|
| rs11041476 | P33241 | LSP1 | miRNATS | Yes | Yes |
| rs11168249 | Q8WUI4 | HDAC7 | TFBS | Yes | Yes |
| rs11676348 | P25025 | CXCR2 | miRNATS | Yes | Yes |
| rs1182188 | Q03113 | GNA12 | TFBS | Yes | Yes |
| rs12103 | Q5TA45 | INTS11 | miRNATS | Yes | Yes |
| rs1598859 | P19838 | NFKB1 | miRNATS | Yes | Yes |
| rs543104 | Q8IZL2 | MAML2 | miRNATS | Yes | Yes |
| rs6087990 | Q9UBC3 | DNMT3B | TFBS | Yes | Yes |
| rs7404095 | P05771 | PRKCB | miRNATS/TFBS | Yes | Yes |
| rs10758669 | O60674 | JAK2 | TFBS | Yes | No |
| rs11130213 | P13798 | APEH | miRNATS | Yes | No |
| rs11130213 | P26927 | MST1 | TFBS | Yes | No |
| rs11130213 | Q9UPA5 | BSN | TFBS | Yes | No |
| rs11209026 | Q5VWK5 | IL23R | miRNATS/TFBS | Yes | No |
| rs11230563 | P30203 | CD6 | miRNATS | Yes | No |
| rs11567699 | P16871 | IL7R | miRNATS/TFBS | Yes | No |
| rs11567701 | P16871 | IL7R | TFBS | Yes | No |
| rs11879191 | Q16543 | CDC37 | miRNATS | Yes | No |
| rs2266959 | P68036 | UBE2L3 | miRNATS/TFBS | Yes | No |
| rs3024495 | P22301 | IL10 | miRNATS/TFBS | Yes | No |
| rs3024505 | P22301 | IL10 | miRNATS/TFBS | Yes | No |
| rs3197999 | P26927 | MST1 | miRNABS | Yes | No |
| rs6667605 | Q92956 | TNFRSF14 | TFBS | Yes | No |
| rs727088 | Q15762 | CD226 | miRNATS | Yes | No |
| rs11229555 | Q6IB77 | GLYAT | miRNATS/TFBS | No | Yes |
| rs13277237 | Q8TAB7 | CCDC26 | miRNATS | No | Yes |
| rs254560 | Q9H5L9 | C5orf66 | miRNATS | No | Yes |
| rs254562 | Q9H5L9 | C5orf66 | miRNATS | No | Yes |
| rs259964 | Q5JPB2 | ZNF831 | miRNATS | No | Yes |
| rs3742130 | Q14330 | GPR18 | miRNATS | No | Yes |
| rs3742130 | Q8NBM4 | UBAC2 | miRNATS | No | Yes |
| rs1131095 | P13798 | APEH | miRNATS | No | No |
| rs12132298 | Q3KP66 | INAVA | miRNATS | No | No |
| rs2382817 | Q8N490 | PNKD | miRNATS/TFBS | No | No |
| rs2382817 | Q969X1 | TMBIM1 | miRNATS/TFBS | No | No |
| rs3749171 | Q9HC97 | GPR35 | miRNATS | No | No |

| | | | | | |
|---|---|---|---|---|---|
| rs41299637 | Q3KP66 | INAVA | TFBS | No | No |
| rs4560096 | Q3MIT2 | PUS10 | miRNATS | No | No |
| rs561722 | Q96DL1 | NXPE2 | miRNATS | No | No |
| rs59655222 | Q3KP66 | INAVA | miRNATS | No | No |
| rs7554511 | Q3KP66 | INAVA | TFBS | No | No |
| rs7657746 | Q2LD37 | KIAA1109 | miRNATS | No | No |
| rs8005161 | Q8IYL9 | GPR65 | TFBS | No | No |
| rs9822268 | P13798 | APEH | miRNATS | No | No |

t
